# EZH2 Abundance Regulated by UHRF1/UBE2L6/UBR4 Ubiquitin System is the Potential Therapeutic Target to Trigger Pigmented Phenotype in Melanoma

**DOI:** 10.1101/2021.03.04.433988

**Authors:** Gamze Kuser Abali, Youfang Zhang, Pacman Szeto, Peinan Zhao, Samar Masoumi Moghaddam, Isobel Leece, Cheng Huang, Jen Cheung, Malaka Ameratunga, Fumihito Noguchi, Miles Andrews, Nicholas C. Wong, Ralf Schittenhelm, Mark Shackleton

## Abstract

Cellular heterogeneity in cancer is linked to disease progression and therapy response, although the mechanisms regulating distinct cellular states within tumours are not well understood. To address this, we identified melanin pigment content as a major source of phenotypic and functional heterogeneity in melanoma and compared RNAseq data from high (HPC) and low pigmented melanoma cells (LPC), revealing the polycomb repressor complex protein, EZH2, as a master regulator of these states. EZH2 protein, but not RNA expression, was found to be upregulated in LPCs and inversely correlated with melanin in pigmented patient melanomas. Surprisingly, conventional EZH2 methyltransferase inhibitors, GSK126 and EPZ6438, had no effect on LPC survival, clonogenicity and pigmentation, despite fully inhibiting methyltransferase activity. In contrast, EZH2 silencing by siRNA strategy or DZNep, MS1943 that reduces EZH2 protein levels, significantly inhibited cell growth in LPCs by hampering ribosome biogenesis. In addition, decline in EZH2 protein level induces pigmented cell phenotype by inducing melanin biosynthesis. Proteasomal inhibitor, MG132 treatment induced EZH2 protein levels in HPCs prompted us to look for differentially regulated ubiquitin system proteins in HPC vs LPCs. UBE2L6, E2 conjugating enzyme has been shown to be downregulated significantly in LPCs by UHRF1-mediated CpG methylation. Both biochemical assays and animal studies demonstrated that UBE2L6 expression decline, in turn, promotes EZH2 protein stability due to lack of ubiquitination on K381 residue in LPCs. UBR4 cooperates with UBE2L6 to facilitate this ubiquitination process. Targeting UHRF1/UBE2L6/UBR4 axis can be a better treatment option to trigger HPC state in melanoma in which conventional EZH2 inhibitors are ineffective.

## INTRODUCTION

The ability of cancer cells to adapt, survive and proliferate during tumor growth, metastasis, and treatment is a major impediment to improving patient outcomes. As the same mechanisms that enable cancer cell adaptation also fuel intratumoral heterogeneity (ITH), understanding drivers of ITH is a high priority in cancer research^1^. Previous reports showed that ITH in melanoma is facilitated by phenotype switching, in which individual cells switch via epigenetic mechanisms between different transcriptional programs, functional states and differentiation phenotypes^2–4^. In melanoma cells, this phenomenon of phenotype switching between the mutually exclusive invasive and proliferative states is under the surveillance of EZH2^3,4^. The proliferative state is characterized by high expression of the melanocyte transcription factor MITF and low Brn2 and Axl expression, while the opposite is true in the invasive state^2,5–10^. Our team has previously shown that melanin pigment content is a major source of cellular ITH in melanoma. To functionally test high (HPC) and low (LPC) pigmented cells from the same melanomas, we developed a novel flow cytometry-based separation method based on the infrared scatter characteristics of different populations within melanomas (Fedele et al., submitted). Strikingly, in cell lines, patient-derived xenografts and fresh patient melanomas, HPCs displayed substantially reduced clonogenicity *in vitro* and tumorigenicity *in vivo*, compared to LPCs (Fedele et al., submitted). Differential gene expression analysis of sorted cell populations by RNA-Seq revealed that HPCs were enriched for melanocytic differentiation gene expression and LPCs with a ribosomal biogenesis gene signature. This indicates critical functional and transcriptional differences in melanoma cells according to their levels of melanin pigment and raises the possibility that the HPC state might be induced for therapeutic benefit.

As a master epigenetic regulator, enhancer of zeste homolog 2 (EZH2) suppresses gene transcription mainly by catalyzing the trimethylation of histone H3 at lysine 27 (H3K27me3). EZH2 plays an important role in maintaining cells in a progenitor-stem cell-like state through silencing genes associated with differentiation^11,12^. EZH2 is aberrantly overexpressed in various malignant tumors, such as prostate cancer^13^, breast cancer^14^, and esophageal cancer^14^, gastric cancer^15^, anaplastic thyroid carcinoma^16^, nasopharyngeal carcinoma^17^, and endometrial carcinoma^18^. EZH2 functions as a critical factor in promoting tumor growth and metastasis in many malignant tumor models^19^.

EZH2 activation by mutations, gene amplification and increased transcription occurred in about 20% of the TCGA cutaneous melanoma (SKCM) cohort^20,21^. In human melanoma clinical samples, EZH2 protein level increases from benign nevi to metastatic melanoma^14,22,23^. EZH2 activation in melanoma represses transcriptional genes associated with tumor suppression, cell differentiation, cell cycle inhibition and repression of metastasis as well as antigen processing and presentation pathways^21,24,25^. It has been known that in melanoma EZH2 expression is upregulated transcriptionally by E2F and c-myc, and it is suppressed by differentiation-promoting factors, such as pRb and p16INK4b^26,27^. MicroRNAs are also capable of downregulating the expression of EZH2 in melanoma^28,29^. However, mechanisms of post-translational EZH2 regulation in melanoma are still largely unknown.

Two recently developed, highly specific EZH2 enzymatic inhibitors, GSK126 and EPZ-6438, are currently in clinical trials for treating lymphomas^19^. Although these EZH2 inhibitors have shown antitumor effects in lymphoma cells with enzyme-activating mutations of EZH2^30–32^ and in ovarian cancer cells with inactivating mutations of ARID1A^33^, certain cancer cells are resistant to the enzymatic inhibition of EZH2 but sensitive to the genetic depletion of EZH2, raising the possibility that EZH2 promotes tumorigenesis via additional mechanisms than its canonical catalytic function. It has been shown that EZH2 enzymatic inhibitors have better inhibitory effect on EZH2 mutant melanoma cells than the wild type^34^. Indeed, independently of its histone methyltransferase activity, EZH2 can promote cancer by stabilizing the PRC2^35^ or by acting as a transcriptional coactivator of androgen receptor^36^, estrogen receptor^37^, β-catenin^37^, and NF-κB^38^. Consequently, targeting EZH2 protein abundance may be more effective than enzymatic EZH2 inhibition for cancers that are dependent on EZH2’s non-catalytic activity.

Cellular protein activity and stability is regulated by post-translational modification^39^. One such modification is ‘ubiquitination’, the reversible addition of ubiquitin protein to the lysine residues of target substrates^40^. This is catalyzed by a series of enzymes: (1) ubiquitin-activating enzymes (E1) use ATP to form ubiquitin-thioester bond; (2) ubiquitin-conjugating enzymes (E2) bind active ubiquitin on their cysteine residues; and (3) ubiquitin-ligase enzymes (E3) interact with E2 enzymes and catalyze the formation of a covalent bond between ubiquitin and its target substrate. E3 ligases regulate substrate specificity. EZH2 protein is subjected to ubiquitin-dependent degradation by several E3 ligases, including β-TrCP, SMURF2, PRAJA1, MDM2 and FBW7 in different cell contexts^41–45^.

Recently, UBE2L6 has been shown as a tumor-suppressor and a novel prognostic marker in melanoma^46^. Here in this study, we identified UBE2L6 as an essential E2 ubiquitin-conjugating enzyme for EZH2, regulating its abundance in melanoma cells and a potential therapeutic target in melanoma to induce HPC state.

## MATERIALS AND METHODS

### Mice

To develop an *in vivo* spontaneous metastasis model, eight-week-old female NOD SCID IL2R−/− mice (NSG) supplied by AMREP, Monash University were injected subcutaneously with empty vector or UBE2LB_pLX307 containing LM-MEL28 B4:F3 melanoma cells (n=11 mice per group). Tumors were measured on a weekly basis using calipers and once the first tumor reached 20mm diameter, the respective mice were sacrificed. All animal experiments were performed in accordance with the approved protocol (number: E/1792/2018/M) by the AMREP Animal Ethics Committee of Monash University.

### Human melanoma tumor samples

19 pigmented and 20 non-pigmented human melanoma tissue sections were obtained from Melanoma Research Victoria (MRV) through the Victorian Cancer Agency Translational Research Program following the approval of the Victorian Government.

### Cell Lines

The HEK293, A375, B16-F10, IGR37 cell lines were from ATCC and cultured under conditions specified by the manufacturer. C006-M1 was from QIMR Berghofer Medical Research Institute, LM-MEL45 was from Ludwig Institute for Cancer Research. LM-MEL28:B4:F3 is the monoclonal line derived from LM-MEL28 cells in our lab previously. Mycoplasma tests were performed in our lab and the short tandem repeat (STR) profiling was done in Australian Genome Research Facility (AGRF) to authenticate the human cell lines.

### Chemicals

The chemicals used for treatment were - GSK126 and EPZ6438 (Selleck Chemicals); MG132 and 5’-Aza-2’-deoxycytidine (Abcam); cycloheximide and DZNep (Sigma) and MS1943 (MedChemExpress). Doses are as indicated in the text or figures.

### Plasmids and siRNA

pCMVHA hEZH2 (#24230), pFlagCMV2-UbcM8 (#12440), pFlagCMV2-UbcM8C86A (#12441), EZH2 E9.3 gRNA (#90684), pCW-Cas9-Blast (#83481), UBE2LB_pLX307 (#98380), pLX307 (#41392), HA-Ubiquitin (#18712), MSCVhygro-F-Ezh2 (#24926) plasmids, and pCMV-VSV-G, pMDL and pRSV-Rev vectors were purchased from Addgene and gift from Marco Herold Lab (WEHI, VIC, Australia), respectively. V5-UBR4-FL was obtained as a gift from Fabio Demontis Lab (St. Jude Children’s Research Hospital, TN, USA)^47^. UBR4 ligand binding domain (aa: 4367-5163) was cloned into FLAG-HA-pcDNA3.1 vector backbone’s NheI/EcoRI restriction sites using the primers hUBR4-LD-NheI-Fw: ATAGCTAGCCCTGGGAACCCGTATAGCAGC and hUBR4-LD-EcoRI-Rv: ATAGAATTCTCAACCGGCCACATCGAGGAACTCTG. pCMVHA hEZH2 vector was used to generate EZH2-K381R mutant vector using the following mutagenesis primers: hEZH2-K381R-Fv: GGAGCTATGATGCTAGATTCCTTGACTCTAAACTCATACAC and hEZH2-K381R-Rv: GTGTATGAGTTTAGAGTCAAGGAATCTAGCATCATAGCTCC with site-directed mutagenesis kit (QuikChange II Site-directed mutagenesis kit, Agilent) following the manufacturer’s instructions. Custom designed siRNA oligonucleotides listed in Table S1 were purchased from Bioneer Pacific. 72 hours after transfection using the Lipofectamine RNAiMAX reagents (Invitrogen) cells were used for functional assays or western blot analysis.

### CRISPR-Cas9-Mediated Gene Editing and Lentiviral transduction

Cells were co-transfected with the puromycin-resistance sgRNA expression vector and the blasticidin-resistance Dox-inducible Cas9 vector. The resistant cells were then selected accordingly and subjected to Western blot analysis for knockout validation. HEK293 cells were co-transfected with pCMV-VSV-G, pMDL, pRSV-Rev and the UBE2LB_pLX307 (Addgene #98380) vector. The respective supernatant was then added to the target cells, followed by the selection of puromycin-resistant cells 48 hours later.

### Co-Immunoprecipitation and HA/ FLAG pulldown assays

1×10^6^ cells were lysed in Co-IP Lysis Buffer (300mM NaCl, 50mM Tris HCL pH 7.4, 0.5% NP40, 0.1% Sodium deoxycholate, 2% SDS) in the presence of cOmplete and PhosSTOP (Roche). Protein-containing supernatants were pre-cleared for 1 h at 4°C with Dynabead® protein G (Thermo Fisher Scientific). Pre-cleared supernatants were incubated overnight at 4°C with 1:1250 dilution of Rabbit (DAIE) mAb IgG isotype control or 1:300 dilution of anti-EZH2 (D2C9) XP Rabbit antibodies. Dynabead® Protein G beads bound to control antibody isotype or desired primary antibody were washed with IP buffer, followed by buffers of increasing salt concentrations (Buffer 1, 50mM Tris, pH 8.0, 150mM NaCl; Buffer 2 50mM Tris, 450mM NaCl and buffer 3, 1M Tris). For Mass Spectroscopy, proteins were eluted by 0.2M Glycine at 4°C and kept in 1M Tris-HCl (pH 8.0) until LC-MS analysis. For HA/FLAG pulldown assays, the cells were lysed in IP Lysis Buffer supplemented with cOmplete protease inhibitor cocktail (Roche), followed by centrifugation at 15,000 rpm at 4°C. Ten percent of the clarified protein solution served as the input control, and 90% was subjected to pulldown analysis through incubation with 25 μl of Pierce anti-HA magnetic beads or Pierce Anti-DYKDDDDK Magnetic Agarose beads (Thermo Fischer Scientific) at RT for 30 minutes. TBST-washed beads were then boiled in 2x SDS-Laemmli Sample Buffer for 10 minutes.

### Western blot

Total protein was extracted from cell lines and tumor xenografts in ice-cold lysis buffer (10mM TRIS-HCL, pH 8.0, 1mM EDTA, 1% Triton X-100, 0.1% Sodium Deoxycholate, 140mM NaCl) in the presence of protease and phosphatase inhibitors after incubation for 1h on ice and centrifugation at 15,000 rpm for 15 min. Proteins were then separated on 4–20% Mini-PROTEAN TGX Stain-Free Protein Gels (BioRad, 4568096), transferred to PVDF membranes followed by incubation with the respective antibodies (Table S1). Signals were detected using Clarity ECL Western blotting Substrate (BioRad) and signal intensities were quantified by ImageJ densitometry analysis software where applicable.

### Quantitative PCR and Methylation-sensitive PCR (MSP)

Total RNA was extracted using Purelink RNA mini isolation kit and Purelink On-Column DNA purification step (Thermo Fischer Scientific) according to the manufacturer’s instructions. Following the synthesis of complementary DNA (cDNA) using SuperScript Vilo cDNA synthesis kit (Thermo Fisher Scientific), quantitative PCR (qPCR) was carried out employing Fast SYBR Green Master Mix (Invitrogen) on a LightCycler 480 Instrument II (Roche). RPLP0 mRNA was included in all qPCR reactions as an internal control. RNA expression changes were determined using a ΔΔCt method^48^. The primers used for EZH2, UBE2L6, and RPLP0 mRNA amplifications are listed in Table S4. To analyze the methylation status of UBE2L6 promoter in DMSO or 5’-Aza treated A375 cells, Quick-DNA Miniprep and the EZ DNA Methylation kit (Zymo Research) were used. The methylated and unmethylated UBE2L6 promoter specific primers (listed in table S1) were designed using the ENCODE CpG island program and MethPrimer software (http://www.urogene.org/cgi-bin/methprimer2/MethPrimer.cgi).

### Cell cycle and apoptosis assays

Cells were fixed and permeabilized with ice-cold 0.5% paraformaldehyde and 70% ethanol, respectively, for 1 h at 4°C. This was followed by staining with propidium iodide (PI) staining solution (1:500 dilution) (Sigma-Aldrich) at 37°C for 30 min and Annexin V-FITC (1:1500 dilution) (Thermo Fischer Scientific) for 15 min at room temperature (RT). Cells were then washed, filtered and subjected to flow cytometry (FACSFusion).

### Cell Proliferation and Clonogenicity assays

To measure cell proliferation rates, equal number of cells were seeded in 6-well plates. Cells were then harvested, stained with trypan blue and counted using a haemocytometer at the desired time points. For assaying clonogenic potential, cells were seeded in 6 cm tissue culture dishes. After 3 weeks of growth, the resulting colonies were fixed with methanol, stained with 0.5% Crystal Violet for 20 min at RT and counted in five random fields using bright field microscopy at 40X magnification. The average colony number per field was then plotted as a function of time.

### Cell invasion assay

Cells were detached by trypsinization and resuspended in serum-free RPMI containing 0.1% BSA and 250,000 cells were plated on top of 100 μl 1:4 diluted growth factor reduced Matrigel in 8.0 μm Boyden chambers and incubated in 600 μl of RPMI medium containing 10% FBS for 48h. Inserts were fixed in 70% EtOH for 15 min and the cells inside of the insert were cleaned by cotton-tipped applicator carefully. Lastly, cells were stained with crystal violet staining solution for 20 minutes at room temperature and rinsed in water by dipping into water-filled container to remove excess CV then air-dried. Cells were counted under inverted microscope in at least 5 random fields and the average number was calculated.

### Cell senescence β-Gal assay

To assess cellular senescence status using the Senescence β-Galactosidase staining kit (Cell Signalling #9860), cells were fixed, washed, then incubated with the β-Galactosidase Staining Solution at 37°C overnight in a dry incubator. Bright field images were taken on a Leica fluorescent microscope for analysis.

### Extracellular melanin assay

For quantitative analysis of extracellular melanin content of B16-F10 cells, the method described by Huberman *et al* was used with some modifications^49^. Briefly, 200 μL of the cell culture medium was transferred to a 96-well plate and the absorbance was read at 405 nm by a spectrophotometer. We used a standard curve obtained from synthetic melanin dissolved in 1 N NH4OH, diluted in culture medium. The absorbance was averaged from three wells and the extracellular melanin content was normalized to the total cell number in the corresponding treatment group.

### Immunofluorescence (IF)

The cells which were treated with either DMSO or 2 μM DZNep for 3 days as well as the FACSorted cells which were allowed to adhere onto slides were first fixed with 4% PFA for IF studies. This was followed by a blocking step using 0.3% Triton X-100 supplemented with 5% normal donkey serum. The samples were then incubated with diluted primary antibody overnight and fluorescent secondary antibody (Table S1). Slides were then washed with PBS, counterstained with DAPI and mounted using Dako Fluorescence Mounting Medium (Dako). Images were then acquired by Leica DM IL LED inverted fluorescent microscope.

### Pigmentation-based flow cytometry sorting of PDX samples

Tumors dissected from euthanized mice were manually dissociated in Hank’s Balanced Salt Solution (without Ca2+ and Mg2+, HBSS−/−), and then processed using the gentleMACS tissue dissociator in Tissue Dissociating media (200 μ/mL Collagenase IV, 5 mM CaCl2 in HBSS −/−). -Following the wash with HBSS−/−, tissues were pelleted at 220g for 4 min at 4°C and then resuspended in 100 units/g of DNase and 5mL/g of warmed trypsin-EDTA. Samples were pelleted again after the addition of equal volumes of cold staining media (0.2% BSA, 10 mM Hepes pH 7.4 (Sigma), 1x PenStrep, Leibovitz’s L15 media (Invitrogen)) and the pellets were then resuspended in cold staining media followed by filtration through a 40 micron filter. An antibody cocktail of anti-mCD31 (endothelial cells), mCD45 (white blood cells), mTER119 (red blood cells) and anti-human HLA-A/B was used to separate the tumoral cells from mouse stroma. In brief, 1X10^6^ cells were incubated in the staining antibody cocktail for 30 min in the presence of single color controls. This was followed by staining with 1:500 dilution of DAPI and 10 μL/mL of DNase. To assess pigmentation, cell sorting was performed on a FACSAria Fusion cytometer using an additional 710nM infrared laser.

### Immunohistochemistry (IHC), Fontana Masson and Schmorl’s staining

Spontaneous metastasis status in *in vivo* models was evaluated by human mitochondrial IHC of liver, lung and lymph nodes^50^. Briefly, slides were de-waxed and antigen retrieval was performed in a pressure cooker at 125°C for 3 min employing an Antigen Retrieval solution (Dako, pH6 for EZH2, pH9 for Anti-human Mitochondrial Antibody). Slides were subsequently incubated with the respective primary and secondary antibodies (Table S3) using an ImmPRESS™ HRP Anti-Mouse IgG (Peroxidase) Polymer Detection Kit (Vector Laboratories) for 60 min at RT. The slides were developed by adding AEC+ High Sensitivity Substrate Chromogen Ready to use (Dako K346111-2). Antibodies used for IHC staining of EZH2 and UBE2L6 in patients’ samples can be found in Table S3. Aperio ImageScope Software was used for evaluation of patients’ IHC and the data were presented either as means of percentages of all immunopositive cells alone or as a classic H-score calculated based on three groups of reaction intensity, i.e. 3+, 2+, and 1+ as follows: H-score: [(3+%cells) x 3 + (2+%cells) x 2 + (1+%cells) x 1]. For EZH2, nuclear reactions were investigated. The five random fields per section were used in all IHC quantitative analyses and the cellular content in each field was comparable within/ among samples. For visualization of melanin in sorted cells (Fontana Masson staining), sorted cells were allowed to adhere on the microscopic slides. The slides were then fixed with 4% PFA washed and incubated in Fontana silver nitrate working solution (2.5% Silver nitrate, 1% ammonium hyrdroxide) at 60°C for 2 hours. A second 2-minute incubation in 0.2% gold chloride solution (Sigma-Aldrich) was performed following several washes. Rinsed slides were then incubated with 5% sodium thiosulfate, counterstained with DAPI and mounted with Dako fluorescence mounting medium (Dako). For Schmorl’s staining, de-waxed sections were rehydrated and stained with Schmorl’s stain (1% Ferric Chloride, 1% Potassium Ferricyanide) for 10 min. Following eosin counterstaining, slides were mounted with DPX Mounting Medium (Thermofisher) for evaluation of melanin content.

### RNA-Seq data and gene set enrichment analysis

FASTQ files were processed using Laxy (https://zenodo.org/record/3767372) which encompasses the RNAsik pipeline (https://joss.theoj.org/papers/10.21105/joss.00583). Briefly, GRCh38 reference genome was used for STAR alignment^51^ and gene expression counts were performed using featureCounts^52^. Gene counts were analysed using Degust (https://zenodo.org/record/3501067) for differential expression analysis. For the Gene Set Enrichment Analysis (GSEA), the gene count matrix for three scr and three siEZH2 samples were uploaded through the desktop software GSEA 4.0.3^53^. To identify significantly enriched gene sets, gene sets from the Molecular signatures database (MSigDB) v7.0^54^ (H: 50 hallmark gene sets; C6: 189 oncogenic signature gene sets) were analyzed using default parameters.

### The Cancer Genome Atlas (TCGA) survival analysis

The clinical data and mRNA expression profiles for skin melanoma samples in TCGA PanCancer Atlas database were accessed through the public cBioPortal (http://www.cbioportal.org).^55^ The high and low expression groups for each gene were defined as above or below the median expression level for the cohort, respectively. Using the R package ‘survminer’ 0.4.4 and ‘survival’ 3.1-11, respectively, the overall survival (OS) Kaplan-Meier curves were plotted and the log-rank test was performed.

### Liquid chromatography–tandem mass spectrometry (LC/MS) interactome and post-translational modification (PTM) analysis

For LC/MS interactome studies, the TCEP (tris(2-carboxyethyl)phosphine)-reduced, alkylated eluents were digested with trypsin at 37°C and the extracted peptides were then cleaned with ZipTips (Agilent) and dried in a SpeedVac (VWR). Using a Dionex UltiMate 3000 RSLCnano system equipped with a Dionex UltiMate 3000 RS autosampler, an Acclaim PepMap RSLC analytical column (75 µm x 50 cm, nanoViper, C18, 2 µm, 100Å; Thermo Scientific) and an Acclaim PepMap 100 trap column (100 µm x 2 cm, nanoViper, C18, 5 µm, 100Å; Thermo Scientific), the tryptic peptides were separated by increasing concentrations of 80% ACN/0.1% formic acid at a flow of 250 nL/min for 90 min and analyzed with a QExactive HF mass spectrometer (Thermo Scientific). The instrument was operated in the data dependent acquisition mode to automatically switch between full scan MS and MS/MS acquisition. Each survey full scan (m/z 375–1575) was acquired in the Orbitrap with 60,000 resolution (at m/z 200) after accumulation of ions to a 3x 10^6^ target value with maximum injection time of 54 ms. Dynamic exclusion was set to 15 seconds. The 12 most intense multiply charged ions (z ≥ 2) were sequentially isolated and fragmented in the collision cell by higher-energy collisional dissociation (HCD) with a fixed injection time of 54 ms, 30,000 resolution and automatic gain control (AGC) target of 2 x 105.

For label-free quantification (LFQ) analysis and protein identification, the raw data files were analyzed using MaxQuant software suite v1.6.5.0 (Tyanova, Temu, & Cox, 2016) against Andromeda search engine (Cox et al., 2011) and in-house standard parameters. The results were analyzed and visualized using LFQ-Analyst (PMID: 31657565).

For PTM analysis, the raw files were searched with Byonic v3.0.0 (ProteinMetrics) using GlyGly at lysine as a variable modification. Only peptides and proteins falling within a false discovery rate (FDR) of 1% based on a decoy database were considered for further analysis.

### Statistical Analysis

Analysis was performed using GraphPad Prism version 7 employing the paired/unpaired Student’s t-test as appropriate.

## RESULTS

### Low pigmented melanoma cells demonstrated an upregulation of EZH2 protein expression with an EZH2-target signature

To decipher the functional differences in melanoma cells according to their levels of melanin deposition, RNAseq was performed on low pigmented cells (LPCs) and high pigmented cells (HPCs) which were originally optimized to be sorted from LM-MEL28: B4:F3 melanoma cells in our lab (Fedele et al., submitted). GSEA analysis between the high- and low-pigmented B4:F3 samples revealed that the LU-EZH2-target-DN gene set was markedly enriched (p=0.006) in LPCs (Fig. 1A and 1B). Next, we compared differentially expressed gene signatures (DEGs) on sorted LPCs and HPCs from B16-F10 cells with those of EZH2 silenced B16-F10 cells measured by RNAseq (Table S1). 31 of 153 LPC upregulated genes were shown to be EZH2-activated genes (Fig. 1C, top panel Venn diagram) and 96 of 209 LPC downregulated genes were identified to be EZH2-repressed genes (Fig 1D, top panel Venn diagram). While the GO biological process analysis of the former 31 activated genes showed that they had enrichment for ribosomal biogenesis (Fig. 1C, bottom panel) the latter 96 repressed genes had significant enrichment for melanin biosynthesis (Fig. 1D, bottom panel). Melanin biosynthesis/melanocytic differentiation genes such as *Oca2*, *Trpm1* and *Met* were over 2-fold upregulated by EZH2 downregulation (Table S2). In addition, *Oca2* is in the top ten genes most negatively correlated with *EZH2* in TCGA skin cutaneous melanoma datasets (p=0.002, Table S3). These data suggested that higher EZH2 abundance/activity in LPCs compared to HPCs might affect melanoma differentiation status by upregulating ribosomal biogenesis and/or downregulating melanin synthesis. To further investigate this, we evaluated the abundance of EZH2 at its mRNA and protein levels, as well as its activity marker, H3K27me3, in HPC and LPC fractions of B4:F3, B16-F10 and C006-M1 pigmented melanoma cell lines. WB and IF staining demonstrated that both EZH2 protein level and H3K27me3 level were higher in the LPCs vs. the HPCs (Fig. 1E-H and Fig. S1A, S1B). Surprisingly, no significant difference was observed between LPC and HPC populations with regard to the EZH2 mRNA levels measured by qRT-PCR (Fig. 1I and 1J and Fig. S1C). A similar inverse association between melanoma cell pigmentation and EZH2 protein level were observed in melanoma cells isolated from *in vivo* PDX melanoma models (Fig. 1K).

**Figure 1.**
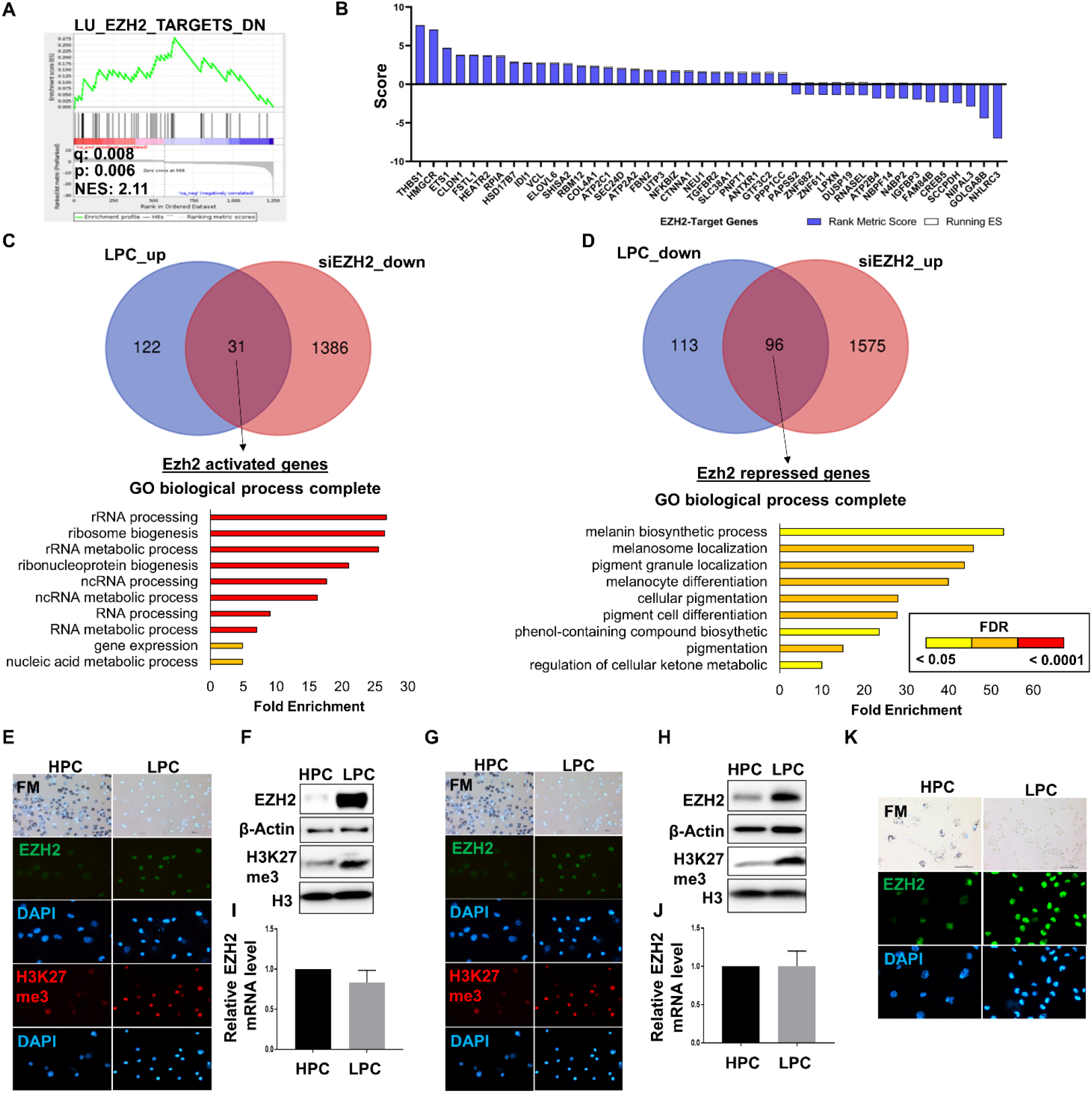
EZH2 protein, but not mRNA level is upregulated in LPCs. (A) Gene set enrichment analysis (GSEA) showing upregulated EZH2 targets in LPCs isolated from LM-MEL-28 B4:F3 cells, and (B) coordinate upregulation of most EZH2-target genes in low-pigmented cells. The ‘Rank Metric Score’ is the signal to noise ratio for each gene used to position the gene in the ranked list. The enrichment score reflects the degree to which the genes included in a gene set are over-represented at the top or bottom of the ranked list of all genes present in the expression dataset. The ‘Running ES’ is the enrichment score for this set at this point in the ranked list of genes. The final enrichment score for a set is the maximum running ES, or the maximum deviation from zero encountered for that set (as illustrated in the lower panel). A positive ES indicates gene set enrichment at the top of the ranked list; a negative ES indicates gene set enrichment at the bottom of the ranked list. (C) Venn diagram showing the overlap of significantly upregulated genes in LPCs of B16-F10 cells relative to HPCs, and of siEZH2 downregulated genes in parental B16-F10 cells relative to scramble control cells (p<0.05, upper panel). GO biological process (complete) enrichment analysis of the 31 common genes showing significantly enriched pathways (p<0.05, lower panel; colour indicates FDR-adjusted p-values). (D) Venn diagram showing the overlap of significantly downregulated genes in LPCs of B16-F10 cells relative to HPCs, and of siEZH2 upregulated genes in parental B16-F10 cells relative to scramble control cells (p<0.05, upper panel). GO biological process (complete) enrichment analysis of the 96 common genes showing significantly enriched pathways (p<0.05, lower panel; colour indicates FDR-adjusted p-values). (E) Bright-field (BF) microscope imaging of Fontana-Masson staining (upper panel) and immunofluorescence (IF) images probed for EZH2 (green) and H3K27me3 (red) in HPCs and LPCs from B4:F3 cells. Nuclear counter-staining is shown by DAPI (blue; lower panels). (F) Endogenous EZH2 and H3K27me3 protein levels measured by western blot and (G) EZH2 mRNA levels were measured by western blot in HPC and LPCs sorted from B4:F3 cells. (G) BF microscope imaging of Fontana-Masson staining (upper panel) and IF images probed for EZH2 (green) and H3K27me3 (red) in HPCs and LPCs from B16-F10 cells. Nuclear counter-staining is shown by DAPI (blue; lower panels). (H) Endogenous EZH2 and H3K27me3 protein levels measured by western blot and (J) EZH2 mRNA levels were measured by western blot in HPCs and LPCs sorted from B16-F10 cells. (K) BF microscope imaging of Fontana-Masson staining (upper panel) and IF images probed for EZH2 (green) and H3K27me3 (red) in HPCs and LPCs from a pigmented melanoma patient-derived xenograft. Nuclear counter-staining is shown by DAPI (blue; lower panels). n=3 biological replicates.

### EZH2 protein level altered pigmentation and oncogenic potential independently of methyltransferase function

To investigate the functional relevance of EZH2 in pigmentation status of melanoma cells, we downregulated EZH2 protein levels and/or its activity in melanoma cells using siRNA or pharmacological inhibitor/degrader approaches. Because high EZH2 expression is only seen in LPCs, the correlation between EZH2 protein levels and pigmentation was next investigated in parental B16-F10 cells which display variable pigmentation levels. Unsorted B16-F10 cells underwent EZH2 silencing using two siRNA constructs. This resulted in a more prominent pigmented phenotype as detected by SSC/NIRSC FACS analysis of melanin content (Fig. 2A), and western blotting showing upregulation of the melanocytic differentiation markers MITF and TYR in EZH2 silenced cells proportionate to the knockdown efficiency (Fig. 2B). EZH2 silencing reduced cell viability in a time-dependent manner (Fig. 2C) and caused an accumulation of cells in the G2/M phase of the cell cycle (Fig. 2D). A more prominent cellular senescence phenotype was also revealed by β-gal staining (Fig. 2E). We found that EZH2 knockdown induced apoptosis, depending on the degree of depletion (Fig. S2A). EZH2 knockout by a CRISPR-Cas9 construct induced MITF levels in pigmented melanoma cell lines (B4:F3, C006-M1 and A375), but this effect was not seen in the non-pigmented SK-MEL-28 cell line (Fig. S2B). An increase in the HPC population frequency was also noted in pigmented melanoma cell lines (data not shown).

**Figure 2.**
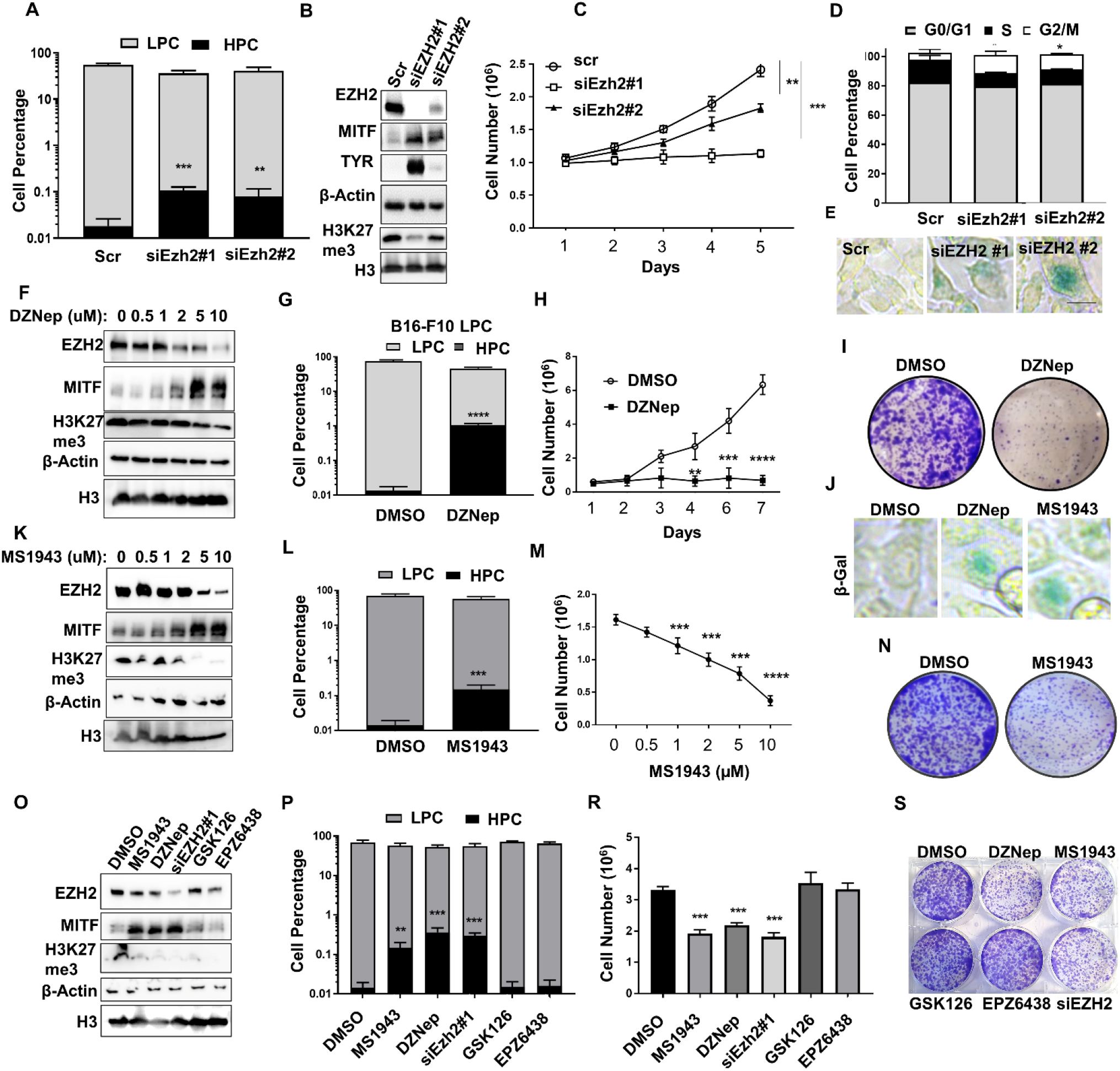
Pharmacological inhibition of EZH2 abundance, but not of its activity triggers the pigmented phenotype in melanoma. (A-E) B16-F10 murine melanoma cells were transfected with either of two siRNAs against Ezh2 or scrambled control, and analysed as follows: (A) HPC and LPC cell percentages were assessed from side-scatter (SSC-A) vs near infra-red scatter (NIRSCA) by flow cytometry; (B) Ezh2, Mitf, Tyr and H3K27me3 protein levels were measured by western blot, including β-actin and H3 as loading controls; (C) growth curves measured by Trypan blue cell counting over 5 days; (D) cell cycle analysis measured by propidium iodide staining; and (E) cell senescence determined by β-gal staining (green). (F-J) B16-F10 cells treated with 2 μM DZNep or DMSO (control) for 3 days were analysed for: (F) Ezh2, Mitf, and H3K27me3 protein level by western blot; (G) HPC and LPC cell percentages by flow cytometry; (H) growth curves over 7 days; (I) clonogenicity after low-density seeding (crystal violet stain); and (J) cell senescence by β-galstaining. (K-N) B16-F10 cells were treated with 2 μM MS1943 or DMSO (control) for 3 days prior to: (K) Western blot analysis of Ezh2, Mitf, and H3K27me3 protein level; (L) HPC and LPC cell percentages by flow cytometry; (M) dose-dependent growth curves; (N) and clonogenicity after low-density seeding (crystal violet stain). (O-R) B16-F10 cells were treated with 2 μM MS1943, 2 μM DZNep, siEzh2 #1, 2 µM GSK126, 2 µM EPZ6438 or DMSO (control) for 3 days prior to: (O) Western blot analysis of Ezh2, Tyr, and H3K27me3 protein level; (P) HPC and LPC cell percentages by flow cytometry; (R) cell numbers counted by Trypan blue; and (S) clonogenicity after low-density seeding (crystal violet stain). Clonogenicity was assessed in pre-treated (3 days) cells seeded at 2000 cells in 6-well plate followed by crystal violet staining (0.5% in methanol) after incubation for 10 days in drug-free media.; Representative images of n=3 biological replicates are shown for western blots (B,F,K,O), clonogenicity plates (I,N,R) and β-gal staining (E,J). * p<0.01, ** p<0.001, *** p<0.0001, **** p<0.00001.

Consistently, when B16-F10 cells were treated with DZNep, a general methyltransferase inhibitor with ubiquitin-mediated proteasomal degradation ability, we found an increase in MITF, a decrease in EZH2, together with a dose-dependent reduction in H3K27me3 through western blotting (Fig. 2F). Meanwhile, we noted a time- and dose-dependent decrease in the macroscopic size of the cell pellet, as well as a more prominent pigmented phenotype at the microscopic level. (Fig. S2C and S2D). This observation was confirmed by SSC/NIRSC FACS analysis showing an increased pigmented cell phenotype of B16-F10 parental and its LPC sorted cells following a 3-day course of treatment with 2 µM DZNep (Fig. 2G and Fig. S2E). Extracellular melanin content was also induced significantly upon EZH2 knockdown or 2 µM DZNep treatment (Fig. S2F). In line with the findings of our siRNA approach, the DZNep-treated cells displayed a markedly reduced cell viability and clonogenicity compared to untreated controls (Fig. 2H and 2I), as well as a significant accumulation of cells in G2/M phases, indicative of cell cycle arrest (Fig. S2G). While apoptosis levels were unaffected, DZNep induced a more senescent phenotype as confirmed by ß-Gal staining (Fig. 1J and Fig. S2J). In addition, we noted a strong upregulation of EZH2 target genes *Cdkn1a* (p21) and *Cdkn2a* (p16) following DZNep treatment of B16-F10 cells (Fig. S2I). We had previously shown in our lab that nucleolar c-Myc and nuclear p53 can serve as LPC and HPC markers, respectively (Fedele *et al*., submitted). DZNep treatment increased nuclear p53 and reduced nucleolar c-Myc levels, indicating an induced HPC differentiation phenotype upon EZH2 inhibition (Fig. S2I). It also reduced ribosome biogenesis as shown by NCL IF (Fig. S2J) which was shown to be upregulated in LPCs (Fedele *et al*., submitted).

EZH2 degradation following treatment with the specific EZH2 degrader MS1943^56^ resulted in simultaneous reduction in EZH2 protein and H3K27me3 levels, as well as an upregulation of MITF levels in a dose-dependent manner (Fig. 2K). Treated cells showed higher melanin content (Fig. 2L), greater senescence, and reduced cell viability and clonogenicity (Fig. 2M, 2N, 2J).

Although EZH2 methyltransferase inhibitors, GSK126 (2 µM) and EPZ6438 (2 µM) fully inhibited EZH2 activity as measured by H3K27me3 levels (Fig. 2O), they had no effect on pigmentation, viability or clonogenicity of B16-F10 cells (Fig. 2P-S).

Overall, we found that specific inhibition of EZH2 methyltransferase activity did not phenocopy perturbations that reduced EZH2 protein level (e.g. siRNA-mediated knockdown, EZH2 degradation by MS1943, or DZNep treatment). This suggests that the EZH2 abundance, but not its methyltransferase activity, may be the key therapeutic target to trigger a more differentiated HPC phenotype, with lower tumorigenic potential, in melanoma.

### EZH2 protein expression is higher in metastases than primary melanomas

While an incremental increase in EZH2 protein level from benign nevi to metastatic melanoma has been observed in IHC data^14,22,23^, our analysis of TCGA melanoma samples showed no correlation between EZH2 mRNA levels and overall patient survival or disease staging in melanoma (Fig. S3A and S3B). The melanoma patient survival rate has also been found to be inversely correlated with EZH2 protein levels^14^. Investigating the EZH2 protein status in our 39 melanoma patients cohort, we found a significant incremental increase in the nuclear EZH2 protein levels from early stage to metastatic melanoma, with metastases containing a higher percentage of LPC than early-stage tumors (Fig. S3C and Fedele *et al*., submitted). Overall, these findings suggest that EZH2 protein, but not its mRNA level, may be a key factor for cellular pigmentation status, melanoma metastasis and patient survival.

### Loss of UBE2L6 reduced ubiquitin-associated proteolysis of EZH2 in LPCs

To understand the discrepancy between melanoma cell pigmentation phenotype and EZH2 mRNA or protein levels, HPCs and LPCs were employed to study the post-translational mechanisms regulating EZH2 protein. Since ubiquitin-mediated proteasomal degradation plays a pivotal role in protein turnover and abundance by affecting protein stability^57^, we assessed EZH2 level pre- and post-treatment with proteasome inhibitor MG132 in LPC/HPC-sorted and non-sorted B16-F10 cells using IF staining and western blot, respectively. As shown in Fig. 3A-3B, MG132 treatment increased the EZH2 protein level in HPCs, but not in LPCs, in a time-dependent manner, suggesting a higher EZH2 turnover in that population.

**Figure 3.**
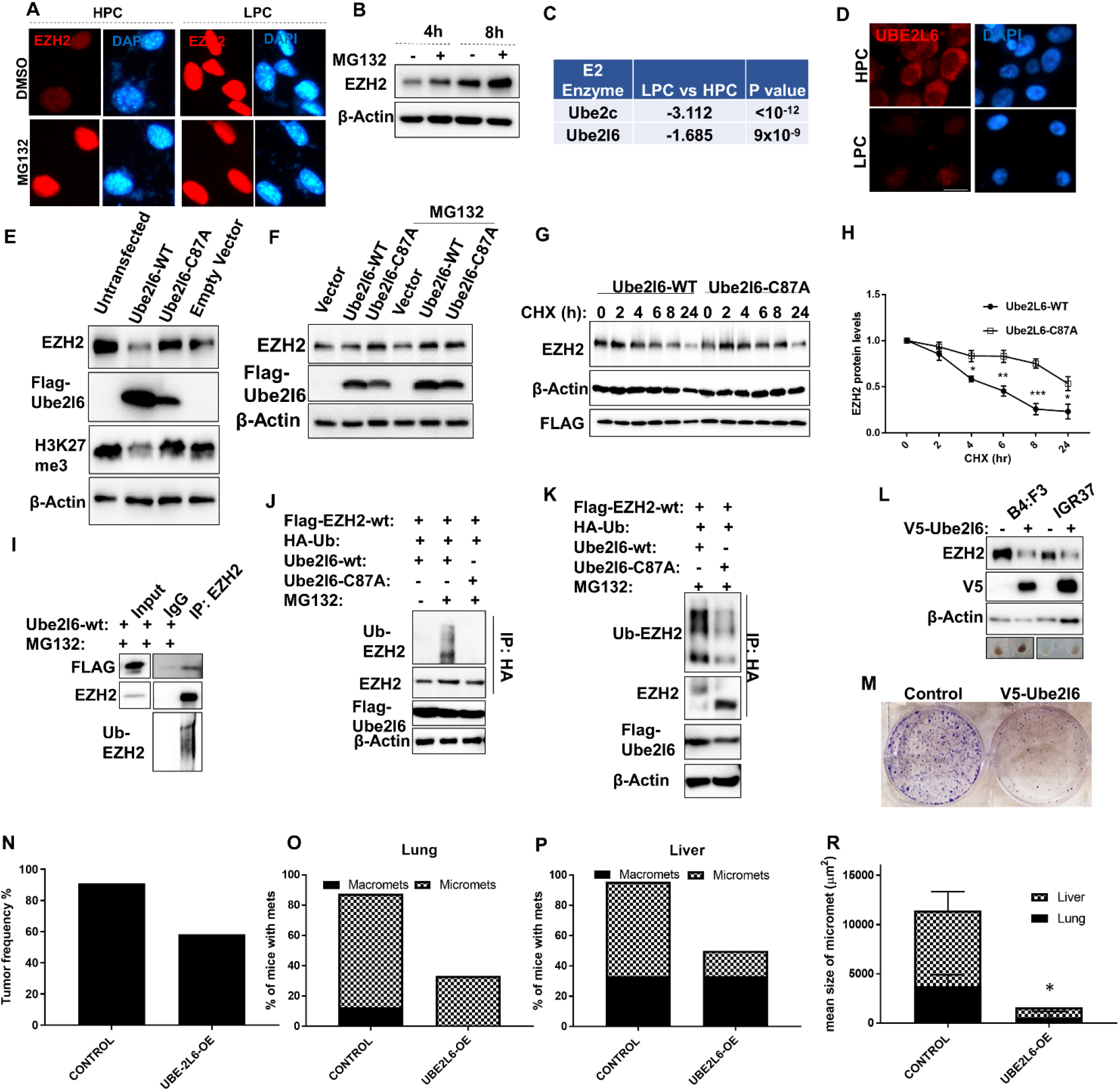
Ezh2 is proteasomally degraded via Ube2l6. (A) EZH2 (red) IF in HPC and LPCs FACS sorted from B16-F10 cells followed by treatment with 10 µM MG132 or DMSO control for 16h. DAPI (blue): nuclei marker. (B) EZH2 protein level was determined by western blot in parental B16-F10 cells treated with 10 µM MG132 or DMSO control for 4h or 8h. (C) E2 ligases that are significantly downregulated in RNAseq data from LPCs vs HPCs. (D) UBE2L6 IF in LPC and HPCs from B16-F10 cells. (E) EZH2 and H3K27me3 levels were determined by Western blot in B16-F10 cells transfected with Flag-tagged Ube2l6-WT, Flag-tagged Ube2l6-C87A (enzyme-dead) or empty vector. (F) As in (E) with or without 10 µM MG132 treatment FOR WHAT DURATION. (G) Stability of endogenous EZH2 protein level was determined by western blot in B16-F10 cells transfected with Flag-tagged Ube2l6-WT or Flag-tagged Ube2l6-C87A followed by 50 µg/mL CHX treatment for the indicated times. (H) Densitometry for western blots shown in (G). n=3 biological replicates. * p<0.01, ** p<0.001, *** p<0.0001. (I) Flag-tagged Ube2l6-WT was overexpressed in B16-F10 cells maintained in the presence 10 µM MG132 for 16h. The interaction between endogenous EZH2 and UBE2L6 was determined by immunoprecipitation with anti-EZH2 antibody followed by western blotting with anti-Flag antibody. (J) HEK293 cells and (K) B16-F10 cells were co-transfected with Flag-tagged EZH2 and Flag-tagged Ube2l6-WT or Flag-tagged Ube2l6-C87A together with HA-tagged ubiquitin in the presence of 10 µM MG132. The ubiquitination of EZH2 was determined by anti-HA IP followed by western blot with anti-EZH2 antibody. (L) B4:F3 and IGR37 cells were infected with V5-tagged empty vector or V5-tagged UBE2L6 lentiviral particles and positive clones were selected by incubation in 2 µg/µL puromycin for 2 weeks. EZH2 level was determined by western blot in the stably-transfected cells and the representative cell pellets were imaged (bottom row). (M) Clonogenicity OF WHAT CELLS assessed by CV staining. (N) Tumor engraftment frequency in NSG mice following subcutaneous injection of B4:F3 cells generated in L. Also shown are the percent of mice with macro- and micro-metastasis to either (O) lung, or (P) liver. (Q) Mean area of micro-metastases in lungs. Control mice number= 11, UBE2L6-OE mice number=11.

To elucidate regulatory mechanisms underlying higher proteasomal degradation of EZH2 in HPCs, we performed a genetic screen for E1-E2 ubiquitination genes to identify the possible candidates involved in EZH2 post-translational regulation. Among E1s and E2s, Uba7 (Fig. S3D), and Ube2l6 and Ube2c (Fig. 3C, Fig., S3E) were unveiled to be significantly decreased at the RNA levels, respectively, in LPCs compared to HPCs. To explore the role of ubiquitination in the fate of EZH2 expression, we focused on UBE2L6 as UBE2C overexpression was found to have a trivial effect on EZH2 protein levels (Fig. S3F) and based on the fact that E1 enzymes have a broader range of substrates. We first confirmed lower expression of Ube2l6 protein by IF staining in LPCs compared to HPCs sorted from B16-F10 (Fig. 3D). Ube2l6-WT overexpression in B16-F10 LPCs dramatically downregulated Ezh2 protein level. On the contrary, cells overexpressing enzyme-dead Ube2l6-C87A showed slightly higher Ezh2 levels as compared to the vector control (Fig. 3E). The Ube2l6-mediated decrease in Ezh2 protein level was reversed by MG132 treatment, pointing out the critical role of proteasomal degradation in the observed effect on Ezh2 (Fig. 3F). Moreover, the inhibitory effect of Ube2l6 on Ezh2 protein level was further verified by Ube2l6 knock-down using the siUbe2l6-3’-UTR construct, which rescued Ezh2 protein expression (Fig. S3G).

EZH2 protein stability was next investigated using cycloheximide (CHX) that inhibits protein synthesis. While EZH2 half-life was found to be around 5h in Ube2l6-WT overexpressed cells, it extended beyond 24h in Ube2l6-C87A overexpressed cells (Fig. 3G, 3H To confirm Ezh2-Ube2l6 protein-protein interaction, EZH2 was immunoprecipitated in Ube2l6-wt overexpressing HEK293 cells treated with MG132, and co-immunoprecipitation of Flag-Ube2l6-wt was observed (Fig. 3I). The presence of higher molecular weight EZH2 bands in western blot (Fig. 3I, lowest panel) prompted us to investigate ubiquitination of EZH2 by UBE2L6. EZH2 was found to be ubiquitinated by UBE2L6-WT, but not by the enzyme-dead form Ube2l6-C87A, in both HEK293 and B16-F10 cell lines (Fig. 3J and 3K). Consistently, DZNep, an inducer of ubiquitin-associated enzymes^41,58^, reduced EZH2 expression in B16-F10 cells and particularly upregulated the UBE2L6 mRNA among the other E2s (>2-fold), further highlighting the role of this enzyme (Fig. S3H, RNAseq data not shown). Ube2l6 knockdown in B16-F10 cells by two different siRNA constructs reversed DZNep-induced EZH2 degradation, TYR expression and cell viability (Fig. S3H). Altogether, UBE2L6 seems to act post-translationally to reduce EZH2 half-life and therefore abundance through ubiquitin-associated proteasomal degradation, thus dampening downstream Ezh2 signalling output in melanoma cells.

### UBE2L6 overexpression diminished EZH2 abundance, tumorigenicity and metastasis *in vivo*

In order to study the functional link between UBE2L6 and EZH2 in melanoma formation, we developed stable UBE2L6-WT-overexpressed pigmented B4:F3 and IGR37 cells. As shown in Fig. 3L-3M and Fig. S3I, S3J, overexpression of UBE2L6-WT in both cell lines dramatically decreased EZH2 protein levels, cell viability and clonogenicity, invasion and visually enhanced cell pigmentation *in vitro*. Following subcutaneous implantation in mice, the frequency of tumor formation was lower in UBE2L6-overexpressing B4:F3 recipient mice (54.5%) as compared to the control group (90%) (Fig. 3N-3O). The difference in tumor volumes in tumor-bearing mice, however, was not statistically significant at 15 weeks. While tumors were partially pigmented in the control mice, they were all pigmented in the UBE2L6-overexpressed group (Fig. S4A), which also demonstrated lower tumors EZH2 protein levels (Fig. S4B) and lower rate of micro and macro metastases to lungs and liver (Fig. 3P, Fig. 3R). Consistent with the primary tumors, macro-metastasesto the liver were low- or partially pigmented in control mice, but pigmented in the Ube2l6-overexpressed group (Fig. S4C). The mean size of micrometastases to lungs and liver were also lower in the Ube2l6-overexpressed group (Fig. 3S and Fig. S4D). On the other hand, all tumor bearing mice developed lymph node metastasis irrespective of Ube2l6 status (data not shown). Taken together, these data demonstrate UBE2L6 as a potential therapeutic target for prevention of melanoma metastasis through EZH2 proteolysis.

### UBE2L6 promoter methylation via UHRF1 resulted in UBE2L6 decline and EZH2 protein upregulation in LPCs

To address whether the regulation of EZH2 by UBE2L6 is relevant in human pigmented melanoma, we performed immunohistochemical staining of these two proteins on 19 pigmented and 20 non-pigmented human melanoma tumor tissue sections. As represented in Fig 4A, tumors with high pigmented cells, defined by Schmorl’s staining, exhibited high UBE2L6, but low EZH2 expression. Conversely, low pigmented cells from the same tumor showed low UBE2L6 and concurrent high EZH2 expression (Fig 4A). Different tumors from pigmented and non-pigmented patient samples with similar stages also showed similar staining pattern (Fig. S5A). We next analysed the UBE2L6 expression in the five random fields of the sections and plotted it against its respective EZH2 protein score. This revealed a significant negative correlation between UBE2L6 and EZH2 protein scores (linear regression R^2^=0.63 and p < 0.0001; Fig 4B) which paralleled the tumor pigmentation status.

**Figure 4.**
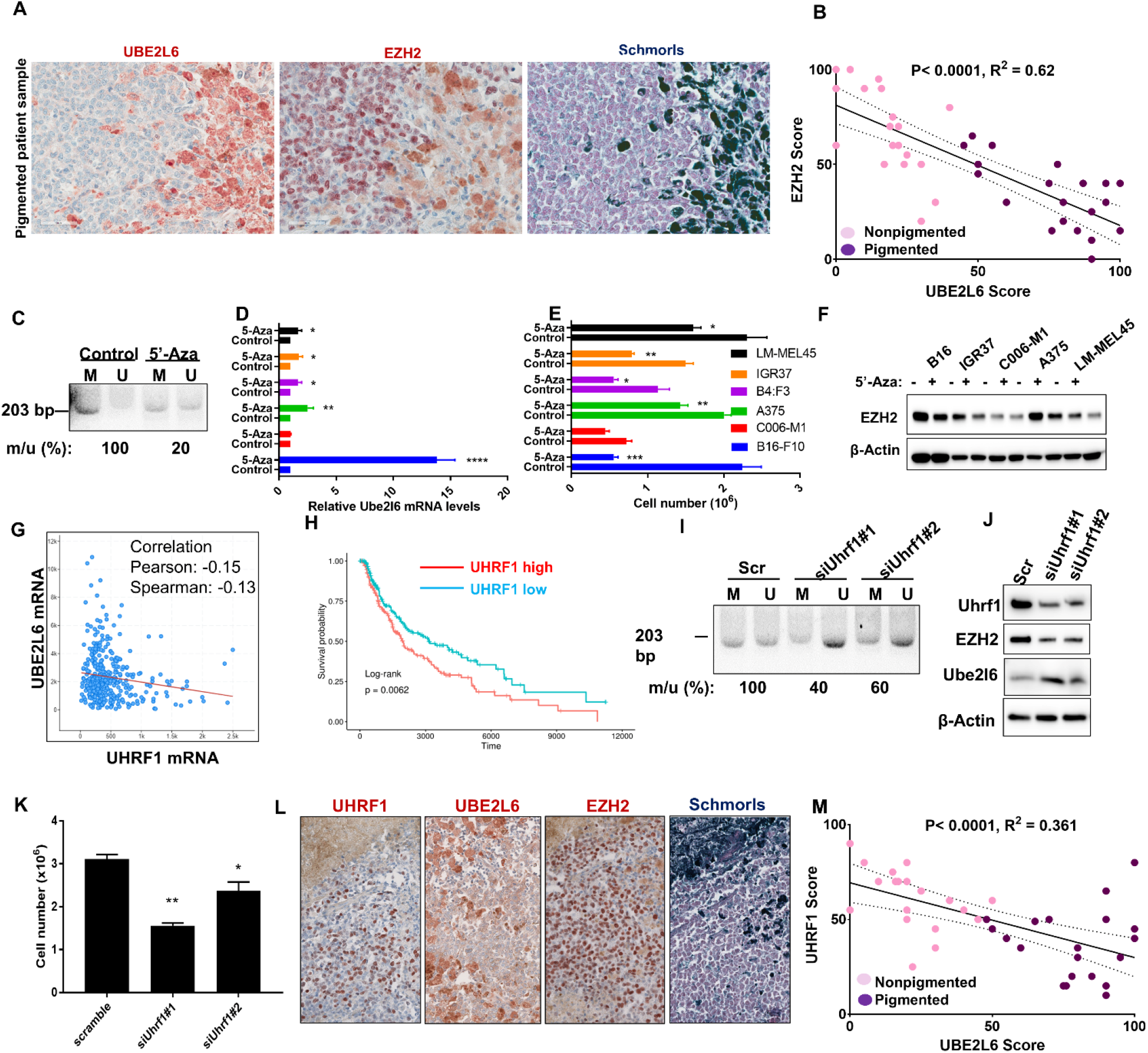
UBE2L6 promoter methylation via UHRF1 results in UBE2L6 decline and EZH2 protein stabilization in LPCs. (A) Immunohistochemical staining of EZH2 and UBE2L6 in representative Schmorl’s stained pigmented human melanoma specimens. Scale bar: 50 μm. (B) UBE2L6 protein scores (x-axis) in melanoma samples negatively correlate with EZH2 protein scores (y-axis) in individual patients. Pink dots correspond to non-pigmented and purple dots to pigmented patient samples. The p value was calculated from a linear regression analysis. R is the correlation coefficient. Protein score = the percentage of immunopositive cells × immunostaining intensity. (C-F) Murine and human melanoma cell lines (as indicated in the panels) were treated with 2 µM 5’-Azacitidine or DMSO (vehicle) for 72h prior to: (C) methylation specific primer (MSP) analysis of the methylation status at the UBE2L6 promoter (M: Methylated specific primer. U: Unmethylated specific primer. The number below the image represents the percent ratio of methylated (m) to unmethylated (u) DNA intensity quantified in ImageJ software); (D) UBE2L6 mRNA quantification with RT-qPCR; (E) viable cell number assessment with Trypan blue cell counting; and (F) EZH2 protein quantification with western blot. (G) Correlation between UBE2L6 mRNA and UHRF1 mRNA levels in TCGA cutaneous melanoma samples. (H) Kaplan-Meier curves of overall survival of TCGA cutaneous melanoma patients (n = xxx patients), stratified by UHRF1 mRNA levels. (I-K) A375 human melanoma cells were transfected with one of two siRNAs against Uhrf1 or scrambled control followed by: (I) MSP analysis of UBE2L6 promoter as in (C); (J) EZH2 and UBE2L6 protein level estimation by western blot; and (K) viable cell number determination by Trypan blue cell counting. (L) Immunohistochemical staining of UHRF1 and UBE2L6 in representative Schmorl’s stained pigmented human melanoma specimens. Scale bar, 50 μm. (M) UHRF1 IF staining of HPC and LPC FACS-sorted from B16-F10 cells. All experiments represent data from n=3 biological replicates. * p< 0.01, ** p< 0.001, *** p< 0.0001.

Next, we analysed TCGA melanoma samples which revealed a correlation between low UBE2L6 expression level and poor melanoma survival (p < 0.0001, Fig. S5B). Investigating the UBE2L6 protein status in our 39 melanoma patient cohort, we found a mild decline (p=0.0575) in the cytosolic UBE2L6 protein levels from low stage to metastatic melanoma which was inversely correlated to the EZH2 protein level (Fig. S5C and Fig. S3C). Interestingly, the UBE2L6 mRNA level was also found to be negatively correlated with its promoter CpG methylation levels (Spearman correlation r = −0.618, Fig. S5D). In view of the above, we hypothesized a possible tumor-suppressor role for UBE2L6 in non-pigmented cells and aimed to evaluate promoter methylation as a mechanism of UBE2L6 regulation in melanoma. To test this, 5-Aza-2′-deoxycytidine (5’-Aza) was used to inhibit DNA methyltransferases (DNMTs) and relieve CpG methylation in A375 cells, followed by methylation-sensitive PCR analysis. The 5’-Aza treatment resulted in an 80% decline in UBE2L6 CpG island methylation, spanning from −582 to −378 relative to TSS (Fig. S5E, Fig. 4C). Moreover, this corresponded to a rise in UBE2L6 expression, a proportional decline in EZH2 protein levels, and a drop in cell viability in B16-F10, A375, LM-MEL28: B4:F3, IGR37 and LM-MEL45 cells, but not in C006-M1 cells (Fig. 4D-F). Thus, UBE2L6 CpG methylation in the promoter can reduce UBE2L6 expression and in turn upregulate EZH2 abundance in non-pigmented cells.

To define the pathway through which UBE2L6 is methylated, we focused on the RING E3 ubiquitin ligase UHRF1 previously shown to downregulate UBE2L6 in cervical cancer cells^59^ and promoted melanoma cell proliferation^60^. An initial analysis of TCGA melanoma samples revealed that Uhrf1 expression inversely correlated with that of UBE2L6 (Spearman correlation r = −0.13, Fig. 4G) as well as melanoma survival (p< 0.0062, Fig. 4H). We next tested whether UHRF1 could regulate UBE2L6 promoter methylation *in vitro*. UHRF1 silencing in A375 cells by two different siRNA constructs resulted in a 40-60% decline in the methylation level of the UBE2L6 promoter depending on the silencing efficacy (Fig. 4I). A concurrent increase in UBE2L6 and decrease in EZH2 protein levels was also seen (Fig. 4J. and Fig. S5F). Cell viability also declined according to the knockdown efficiency in different melanoma cell lines (Fig. 4K and Fig. S5G). IHC staining of UHRF1 in pigmented human melanoma samples also showed that highly pigmented cells exhibiting higher UBE2L6 and lower EZH2 expression levels, also had lower UHRF1 expression (Fig. 4L and 4M, linear regression R^2^=0.52 and p< 0.0001). In summary, these data clearly demonstrated that UBE2L6 promoter methylation via UHRF1 could result in UBE2L6 decline and in turn, EZH2 stabilization in LPCs.

### E3 ligase UBR4 interacts with UBE2L6 to ubiquitinate the K381 residue of EZH2 protein

It has been shown that the activity of E2 conjugating enzymes relies on their cooperation with an E3 ligase through which direct interaction with their substrates happens. To identify the possible E3 ligase(s) interacting with EZH2, EZH2 co-immunoprecipitated lysates obtained from a variety of melanoma cell lines, including A375, LM-MEL28:B4:F3, B16-F10, LM-MEL45 and IGR37 were subjected to LC-MS to investigate EZH2 interaction networks. Among the interacting proteins, UBR4 was the only common E3 ligase identified (Fig. 5A). However, no significant difference in the UBR4 expression level was seen between HPCs and LPCs (Fig. S5H) and pigmented patient samples (Fig. S5I) contrary to our UBE2L6 results (Fig. 3D). To confirm the possibility of UBR4 as a potential EZH2 E3 ligase, the interaction between EZH2 and UBR4-LD (Ligase Domain) was assessed by CoIP coupled western blotting. EZH2 was shown not only to interact with the E3 ligase domain of UBR4, but also its interaction was enhanced by MG132 treatment (Fig. 5C and Fig. S5J). Furthermore, EZH2 protein level was diminished by UBR4-LD and UBR4-FL (Full-Length) overexpression in HEK293 cells (Fig. 5D). As expected, UBR4 knockdown in LM-MEL28:B4:F3, C006 and HEK293 cells increased EZH2 protein levels (Fig. 5E) and its stability (Fig. 5F) via decreased EZH2 ubiquitination as measured by *in vitro* ubiquitination assay (Fig. 5G) and ubiquitin (Ub) specific antibody binding (Fig. 5H). UBR4-FL overexpression, on the other hand, increased EZH2 ubiquitination (Fig. 5H). Taken together, these results indicated that UBR4 is an EZH2 E3 ligase.

**Figure 5.**
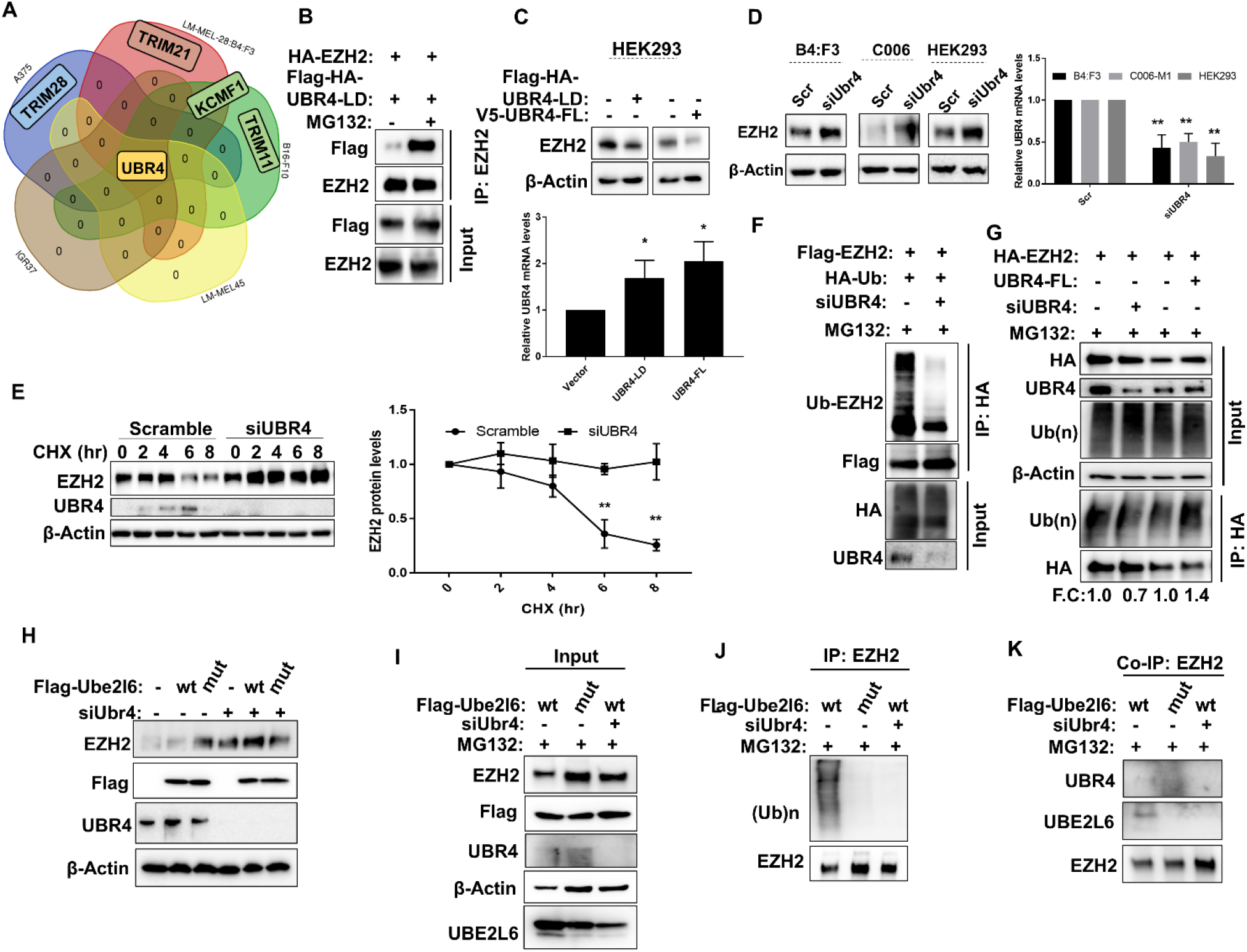
UBR4 is the E3 ligase of EZH2. (A) Venn diagram depicting the overlap of proteins co-immunoprecipitating with EZH2 from C006-M1, LM-MEL-28:B4:F3, IGR37, A375 and LM-MEL-45 melanoma cells (all data derived from n=3 biological replicates). (B) HA-tagged EZH2 and Flag-HA-tagged UBR-LD were co-expressed in HEK293 cells maintained in the presence or absence of MG132. The interaction between EZH2 and UBR4-LD was determined by immunoprecipitation with anti-EZH2 antibody followed by western blotting with anti-Flag antibody. (C) EZH2 protein level was determined in HEK293 cells overexpressing either UBR4-LD or UBR4-FL. UBR4 overexpression efficiency was measured by qPCR (lower panel; data show quantification relative to vector control, error bars indicate +SD). (D) UBR4 in B4:F3, C006-M1 and HEK293 cells was knocked down by siRNA, and EZH2 level determined by western blot (left panel) and the knockdown efficiency of UBR4 was measured by qPCR as in (C) (right panel). (E) Stability of EZH2 protein level under different UBR4 conditions (unaltered vs knockdown) was determined by western blot. Representative blot shown at left, time-course plot at right. (F) The ubiquitination of EZH2 was determined by anti-HA IP followed by western blot. (G) HEK293 cells with or without siUBR4 were transfected with HA-tagged EZH2 or V5-tagged UBR4-FL followed by 50 µM MG132 treatment for 4h. The ubiquitination of EZH2 was determined by western blot with anti-ubiquitin (Ub) antibody. (H) B16-F10 cells with or without siUBR4 were transfected with either Flag-tagged Ube2l6-WT (“wt”) or Flag-tagged Ube2l6-C87A (“mut”) and EZH2 level determined by western blot. (I-K) To evaluate the EZH2/UBR4/UBE2L6 interaction, B16-F10 cells with or without siUBR4 were transfected with either Flag-tagged Ube2l6-WT (“wt”) or Flag-tagged Ube2l6-C87A (“mut”) followed by 50 µM MG132 treatment for 4h and then: (I) EZH2, UBR4 and UBE2L6 protein estimation by western blot; (J) determination of ubiquitination of EZH2 by western blot with anti-Ub antibody; and (K) co-immunoprecipitation of UBR4 and UBE2L6 with EZH2 antibody. All data represent n=3 biological replicates. ** p< 0.001.

To outline the possible cooperative interplay between UBE2L6/UBR4 and EZH2, Ube2l6-WT or Ube2l6-C87A overexpressing B16-F10 cells were subjected to Ubr4 silencing. Interestingly, Ubr4 silencing reversed Ube2l6-WT-mediated ezh2 degradation (Fig. 5I) and ubiquitination in B16-F10 cells (Fig. 5J and 5K). Furthermore, an interaction between endogenous Ubr4 and Ezh2 was demonstrated by Ube2l6-WT overexpression, but not by the Ube2l6-C87A mutant (Fig. 5L). More importantly, the UBE2L6:EZH2 interaction was ablated upon UBR4 silencing (Fig. 5L). These data strongly support the fact that UBE2L6 and UBR4 bind together to form a complex with EZH2 to facilitate EZH2 ubiquitination in melanoma cells.

Next, we aimed to determine the ubiquitination site on EZH2 through which the UBE2L6/UBR4 ubiquitin-enzyme system interacts. Although EZH2 ubiquitination was predicted to be at K381 (human)/ K376 (mouse) at the highest confidence *in silico* by the UbPred program^61^ (Fig. 6A), endogenous ubiquitinated EZH2 was undetectable by LC-MS in all tested melanoma cells in which UBE2L6 levels were extremely downregulated (data not shown). When we assessed Ube2l6-WT overexpressing B16-F10 cells, K376 ubiquitination on EZH2 was detected by LC-MS analysis (Fig. 6B). However, LC-MS could not detect a ubiquitination site upon Ube2l6-C87A overexpression or Ubr4 silencing together with Ube2l6-WT overexpression in order to verify the cooperative interplay between UBE2L6 and UBR4 for EZH2 K376 ubiquitination. For confirmation purposes, we then performed a sequence alignment analysis which revealed that the K381 residue is conserved between *Homo sapiens* and *Caenorhabditis elegans*, thus pointing out the critical role of this lysine residue in EZH2 regulation (Fig. 6C). Comparing K381 mutant and wild-type EZH2 in HEK293 cells, the K381R mutant form showed dramatically enhanced protein stability compared with wild-type EZH2 (Fig 6D) and the mutant EZH2 protein was not subjected to UBE2L6-mediated degradation in B16-F10, A375 and IGR37 cells (Fig. 6E and 6F). In turn, cell viability and invasiveness were restored by the overexpression of wild-type EZH2 and exceeded by the overexpression of EZH2-K381R mutant protein in UBE2L6 stably overexpressing cells (Fig. 6G, H). These data clearly indicate that UBE2L6 acts as a tumor suppressor in melanoma cells by coupling with ubiquitin E3 ligase UBR4 for K381 (human) residue-directed ubiquitin mediated degradation of EZH2.

**Figure 6.**
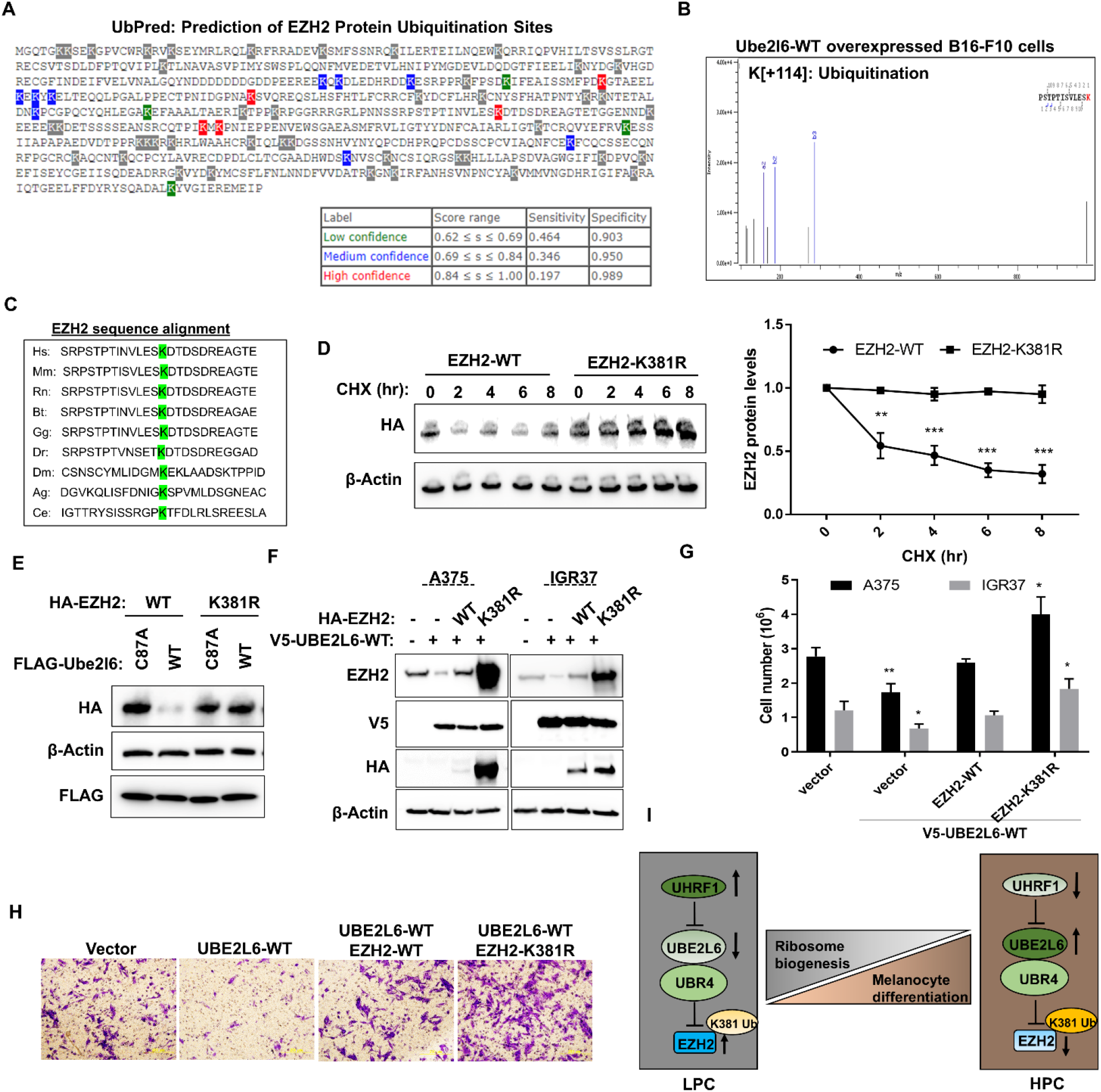
K381 residue on EZH2 is ubiquitinated by UBE2L6/UBR4 in melanoma cells. (A) Prediction of ubiquitination sites on human EZH2 was made using the UbPred program showing low (green), medium (blue) and high (red) confidence lysine ubiquitination sites (www.ubpred.org) (B) LC-MS analysis of mouse EZH2 K376 ubiquitination spectrum (C) Sequence alignment of residues 368-392 of human EZH2 protein against Mus musculus (mouse), Rattus norvegicus (rat), Bos taurus (cow), Gallus gallus (chicken), Danio rerio (zebrafish), Drosophila melanogaster (fruit fly), Anopheles gambiae (mosquito), Caenorhabditis elegans (worm) showing conservation of the K381 residue (D) HEK293 cells with HA-tagged EH2-WT or HA-tagged EZH2-K381A were treated with 50 µg/mL CHX for the durations indicated. Stability of HA-tagged EZH2 protein level was determined by western blot with anti-HA antibody. Representative blot (left) and time course of protein level (right) are shown. (E) B16-F10 cells were transfected with either Flag-tagged Ube2l6-WT or Flag-tagged Ube2l6-C87A (mutant) and HA-tagged EZH2-WT or HA-tagged EZH2-K381A and HA-EZH2 level was determined by western blot with anti-HA antibody. (F) A375 and IGR37 cells transduced with V5-UBE2L6-WT were transfected transiently with either HA-tagged EZH2-WT or HA-tagged EZH2-K381R. Endogenous EZH2 level was determined with EZH2 antibody, HA-EZH2 level with anti-HA antibody and V5-UBE2L6 level with V5 antibody 48h post transfection. (G) Cell viability was determined by Trypan blue cell counting 48h post transfection. (H) Invasion of cells treated as in (F) was performed 72h post transfection using Boyden chamber matrigel invasion assays followed by crystal violet staining. (I) Proposed model. All data represent n=3 biological replicates. * p< 0.01, ** p< 0.001, *** p< 0.0001.

In summary, we identified that UBE2L6 promoter hyper-methylation via UHRF1 results in a decline in UBE2L6 expression in LPCs which in turn upregulates EZH2 protein stability due to the lack of ubiquitination on its K381 residue. Overall, perturbations which decrease EZH2 protein stability/abundance may promote a phenotypic switch from LPCs to HPCs, associated with a less tumorigenic and more senescent status (Fig. 6I).

## DISCUSSION

Our previous experiments showed that widespread heterogeneity in melanin pigment content identifies cells of markedly different tumor forming and sustaining potential. In cell lines, patient-derived xenografts and fresh patient melanomas, HPCs displayed substantially reduced clonogenicity in vitro and tumorigenicity in vivo, compared to LPCs. Through conditional gene expression analysis, we identified a low pigment associated transcriptional signature of EZH2 that represses melanocytic differentiation genes and upregulates ribosome biogenesis regulatory genes. Here, we identified EZH2 as a master regulator of melanoma ITH and promoter of tumorigenic melanoma cell states. Importantly, genetic or pharmacological depletion of EZH2, but not simply inhibition of its methyltransferase activity, inhibits melanoma cell tumorigenicity and induces a HPC state, indicating a critical methyltransferase-independent function of EZH2 in melanomagenesis. These novel functions are subject to UHRF1/UBE2L6/UBR4-mediated degradation of EZH2, which may be a practical strategy to trigger the less aggressive HPC state in melanoma. This study also identifies not only EZH2, but also UHRF1 and UBE2L6, as novel markers of melanoma ITH and oncogenic potential.

Our data elucidate the methylation-independent regulation of clinically relevant melanoma functions by Ezh2. Previous studies implicate genes involved in melanin biosynthesis and melanocytic differentiation – including *Mitf*, *Tyr*, *Tyrp1*, *Dct*, *Mlana*, *Pmel* and *Oca2* – as key targets of Ezh2-directed histone methylation^24,25^, leading to altered pigmentation and differentiation status^62^. Indeed, we found that Ezh2 protein level was inversely associated with pigmentation in both sorted melanoma cells and in clinical samples but selective Ezh2 methyltransferase inhibition using GSK126 or EPZ6438 was not altered melanocytic differentiation gene expressions and pigmentation. While off-target drug effects at the higher doses used in other studies may explain this discrepancy, we demonstrated that biologically relevant effects of Ezh2 manipulation are not dependent on its methyltransferase activity. Specifically, Ezh2 enzymatic inhibitors have less effect on clonogenicity and metastasis in vivo (Kuser-Abali et al., manuscript in preparation) than Ezh2 depletion by either genetic manipulation or pharmacological degradation using DZNep or MS1943. In this regard, our data are consistent with a previous study showing a significant reduction of tumor volume and lymph node metastasis, and absence of lung metastasis in an Ezh2 conditional knockout mouse model with NrasQ61K mutant background28. Relative to Ezh2 methyltransferase inhibition using GSK503, knockout mice showed less metastasis and better survival^24^. Our data extend prior work showing that Ezh2 knockdown reduced melanoma proliferation, induced a senescent phenotype, and inhibited the growth of xenografts in mice^23^, by revealing that Ezh2 has methyltransferase independent, non-catalytic function(s) impinging on melanoma pigmentation, tumorigenesis and invasion.

In melanoma, EZH2 may act at a specific step in the multi-stage metastatic process. It is known that phenotype switching produces melanoma cell populations of mixed tumorigenic potential. Invasive melanoma cells have lower pigment levels compared with non-invasive cells^4,7,62^ and Ezh2 promotes the low-pigment phenotype in vivo by suppressing Oca2, likely among other targets^62^. We demonstrated that control melanoma cells (LM-MEL28:B4:F3) mostly formed partially pigmented tumors (high EZH2 protein levels) following subcutaneous implantation, with non-pigmented metastatic foci in the liver. On the other hand, UBE2L6 over-expressing melanoma cells (low EZH2 protein levels) formed highly pigmented tumors with either no metastatic foci or pigmented foci of a smaller size. Pinner et al.,^7^ showed that non-pigmented invasive melanoma cells metastasize to lung and lymph node, but reverted to a pigmented phenotype after colonization; we did not observe this phenotype switch in our study. Similarly, it was suggested that EZH2 is required for efficient lung and lymph node colonization by metastatic melanoma^24,62^. Consistent with this data we found that metastasis into lung and liver were reduced by UBE2L6 overexpression in mice. However, all tumor-bearing mice developed lymph node metastasis regardless of UBE2L6 status. These discrepancies may arise from differences in melanoma cells used, host mouse models, and experimental duration between studies. EZH2 may be involved not in the early dissemination into lymphoid organs but in subsequent targeting and colonization of visceral (lung/liver) sites. Alternatively, it is possible that other UBE2L6 downstream targets may be critical regulators of melanoma metastasis. However, we have shown that cell viability and invasiveness were restored by the overexpression of wild-type EZH2 and exceeded by the overexpression of EZH2-K381R mutant protein in UBE2L6 stably overexpressing melanoma cells *in vitro*.

The tumor-suppressive/anti-metastatic role of UBE2L6 in vivo has not previously been described and is mediated by the ubiquitin conjugating function of UBE2L6 in melanoma working cooperatively with the E3 ligase UBR4, leading to EZH2 protein degradation. Ubiquitin-mediated proteasomal degradation is critical for efficient protein turnover by reducing the protein stability, thereby protein level^57^. UBE2L6 is also the E2 conjugating enzyme for Interferon Stimulated Gene 15 (ISG15), a 15 kDa ubiquitin-like protein modifier which can be conjugated to protein substrates in order to modify their functions. However, we did not see an effect of UBE2L6 on EZH2 ISGylation in melanoma (data not shown). UBE2L6 abundance is itself regulated by UHRF1, which induces suppressive promoter hyper-methylation of UBE2L6 in LPCs and thus exhibits competing effects on downstream EZH2 protein level. UHRF1 acts as an oncogene in various cancers, including melanoma^59,60^ and has been shown to downregulate UBE2L6 in cervical cancer^59^. In melanoma patient samples, EZH2 protein level was enhanced and UBE2L6 protein level slightly reduced in metastatic lesions (Stage III/IV) in which LPC percentage is higher than in primary tumors (Stage I/II) (Fedele et al., submitted). Thus, loss of UHRF1/UBE2L6/UBR4 mediated EZH2 degradation in LPCs, and enrichment of the LPC phenotype, may explain the higher EZH2 protein levels seen in higher-stage melanomas.

Revealing the molecular events involved in methyltransferase independent Ezh2 function opens new therapeutic directions. Previous reports showed that DZNep reduces both EZH2 protein stability and (consequently) activity by upregulating the E3 ligase PRAJA1 in breast cancer^41^. In this study, we revealed that DZNep reduces EZH2 protein stability by inducing UBE2L6 expression in melanoma cells. Although DZNep is effectively anti-tumorigenic, due to its short half-life (1.1 h in rats and 0.9 h in mice)^63,64^ and broad effect on the ubiquitin system, anti-tumor studies using DZNep have not progressed beyond the pre-clinical phase. However, destabilization of EZH2 by targeting its ubiquitination enzymes may offer an alternative therapeutic approach to treat melanoma. Furthermore, as UBE2L6 and UHRF1 protein are differentially abundant between LPCs and HPCs, these may be useful biomarkers of cellular phenotype and aggressive potential.

## Acknowledgement

The FACS experiments were carried out in the AMREPFlow core flow cytometry facility and the LC-MS experiments were done in the Monash Proteomics and Metabolomics facility. Melanoma patient sample blocks were provided by the Melanoma Research Victoria through the Victorian Cancer Agency Translational Research Program.

## Supplementary Figure Legends

**Figure S1.**
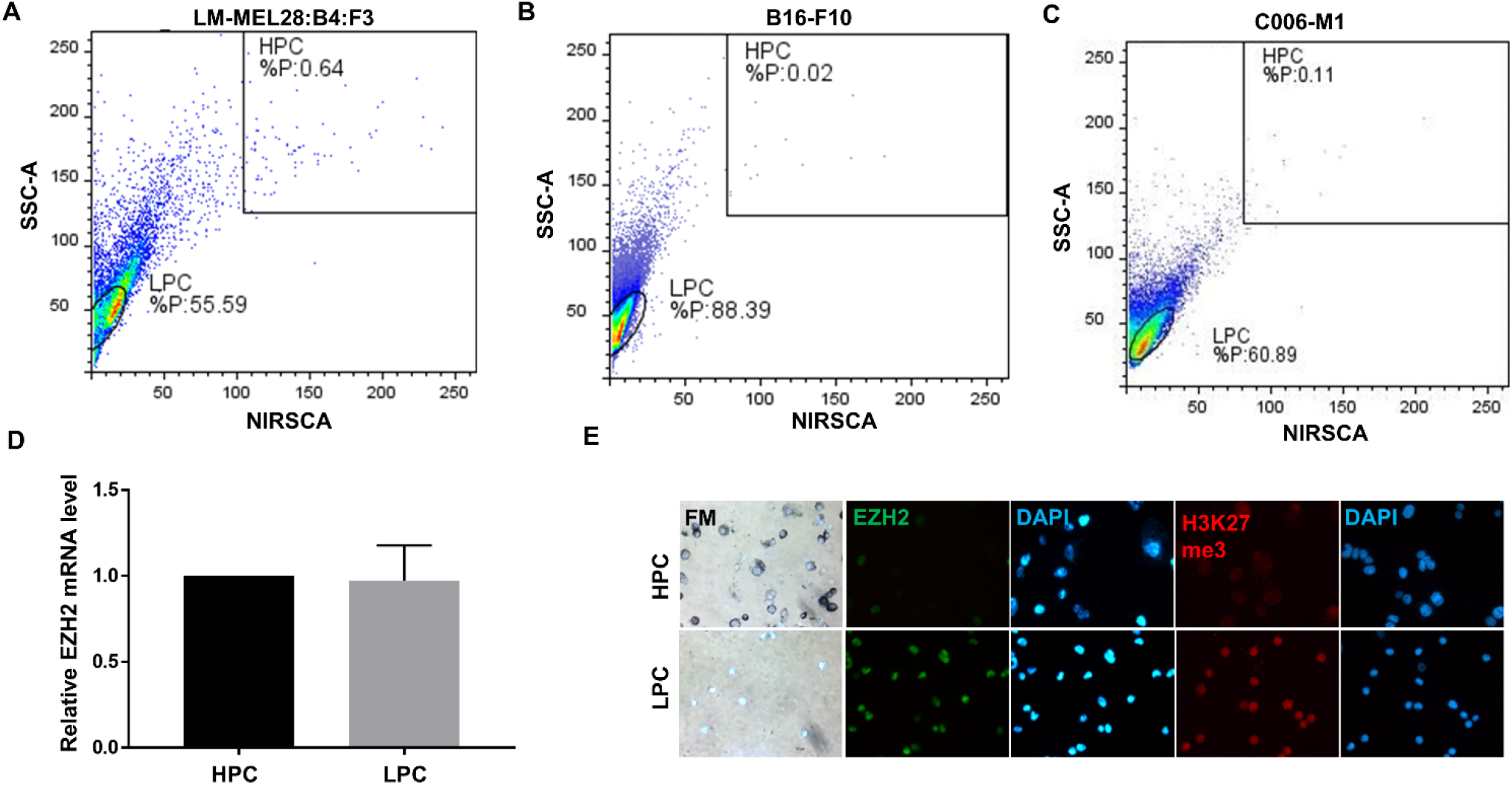
EZH2 protein, but not mRNA level is upregulated in LPCs from C006-M1 cells. (A) SSCA vs NIRSCA FACS analysis of LM-MEL24:B4:F3, B16-F10 and C006-M1 cells (B) EZH2 qRT-PCR of HPC and LPCs from C006-M1 cells. n=3 biological replicates. (C) Bright-field (BF) microscope imaging of Fontana-Masson staining (upper panel) and immunofluorescence (IF) images probed for EZH2 (green) and H3K27me3 (red) in HPCs and LPCs from C006-M1 cells. Nuclear counter-staining is shown by DAPI (blue; lower panels).

**Figure S2.**
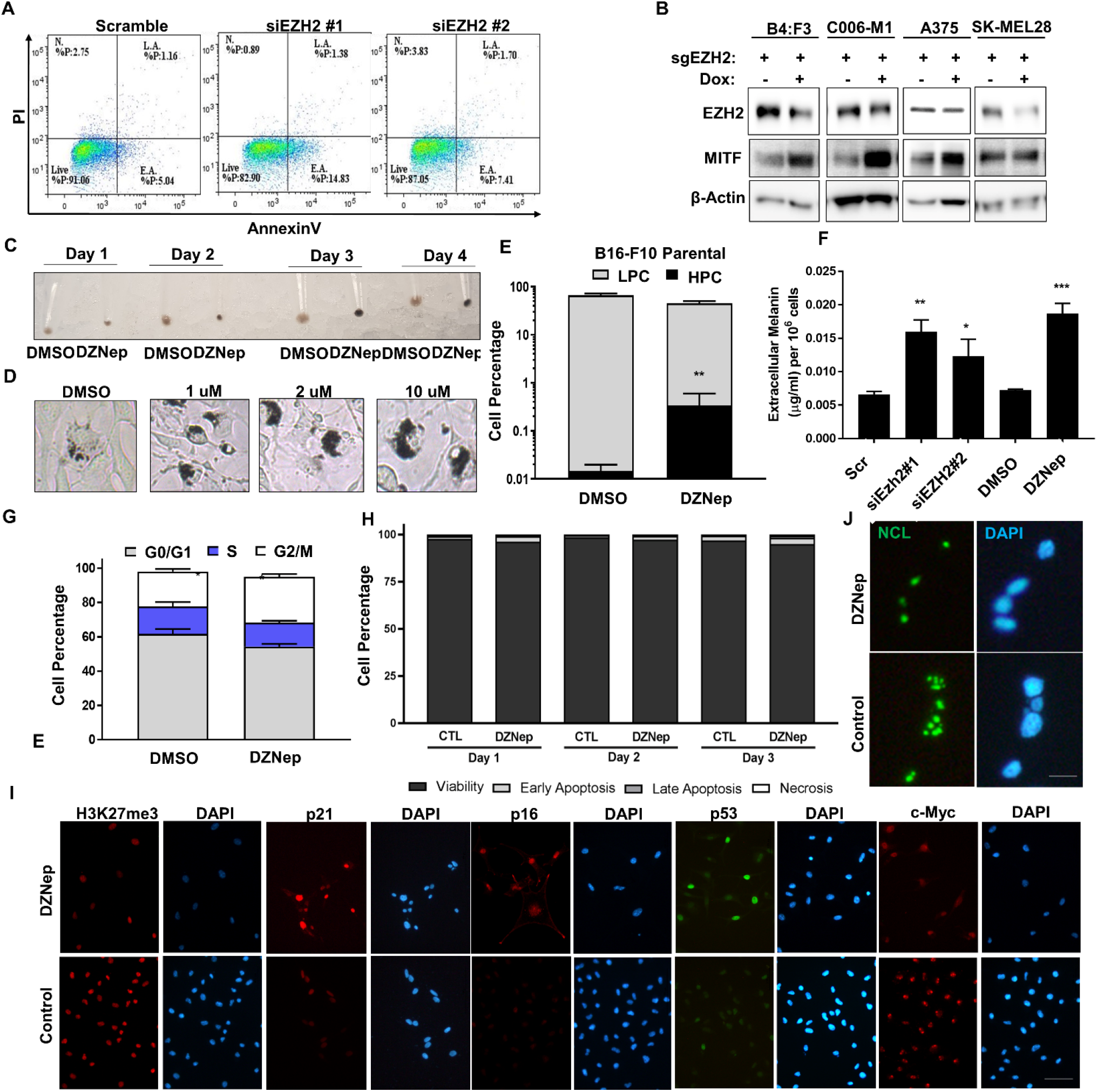
Manipulation of EZH2 protein levels by DZNep or knockout switches HPC phenotype. (A) AnnexinV/PI apoptosis analysis in B16-F10 cells transfected with EZH2 siRNA#1, #2 or scramble control for 3 days. (B) Western blot analysis of EZH2, and MITF in control, EZH2-knockout B4:F3, C006-M1, A375 and SK-MEL28 cells. (C) Cell pellet imaging of B16-F10 cells treated with DMSO or 2 µM DZNep for the indicated times. (D) Bright-field microscopy imaging of B16-F10 cells treated with DMSO or DZNep for 3 days in a dose-dependent manner. (E) HPC and LPC cell percentage analysis in B16-F10 cells treated with DMSO or 2 µM DZNep for 2 days. (F) Extracellular melanin levels were determined by melanin assay in B16-F10 cells transfected with scramble control or siEZH2#1 and #2 or treated with DMSO or 2 µM DZNep for 3 days. (G) Cell cycle analysis by PI analysis in B16-F10 cells treated with DMSO or 2 µM DZNep for 3 days and (H) AnnexinV/PI analysis for the indicated times. (I) p21, p16, p53 and c-Myc IF staining in B16-F10 cells treated with DMSO or 2 µM DZNep for 3 days. n=3 biological replicates. * p< 0.01, ** p< 0.001, *** p< 0.0001.

**Figure S3.**
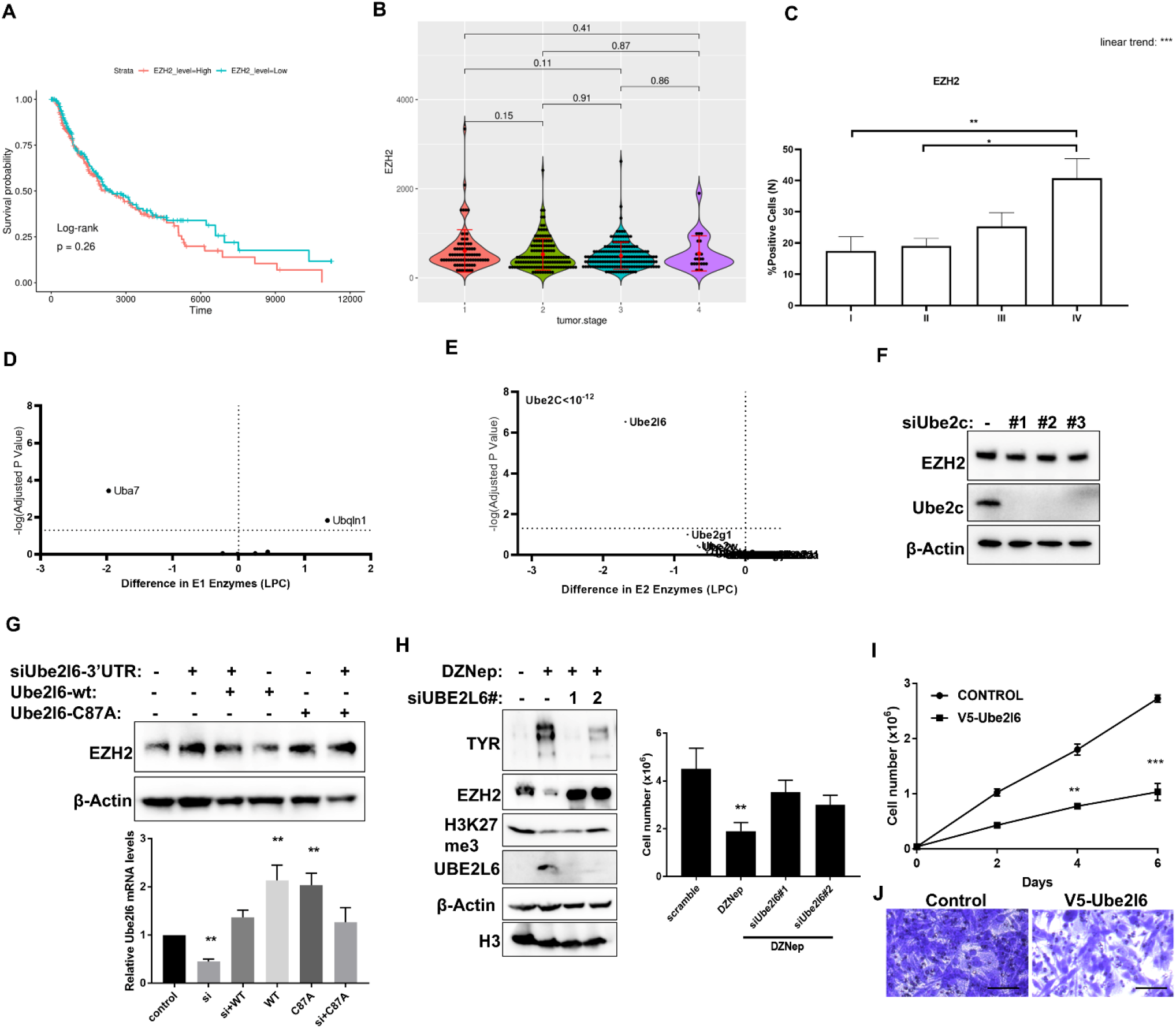
EZH2 protein, but not mRNA correlates with melanoma survival and staging and Ube2l6 is the E2 enzyme of EZH2. (A) Kaplan-Meier survival and (B) tumor staging analysis of TCGA cutaneous melanoma patients (n = 427 patients), stratified by EZH2 mRNA levels. (C) Melanoma tumor staging in our human melanoma patient cohort stratified by nuclear EZH2 protein levels. Protein score = the percentage of immune-positive cells (D) Genetic candidate expression screen of E1 ubiquitination genes from RNAseq data on sorted LPCs and HPCs from B16-F10 (E) Western blot analysis of EZH2 and Ube2c in B16 cells transfected with scramble or three different siUbe2c oligos (F) Endogenous EZH2 protein levels were measured by western blot in B16 cells transfected with Ube2l6-WT or Ube2l6-C87A with or without siUbe2l6-3’UTR oligos (G) Western blot analysis of EZH2 and TYR in B16 cells with or without two different siUbe2l6 oligos treated with 2 μM DZNep for 72h. (H) Cell viability was measured for 6 days by trypan blue counting and (J) boyden chamber matrigel invasion assay in B4:F3 cells that were infected with V5-tagged empty vector of V5-tagged UBE2L6 lentiviral particles. Scale bars = 200 µm n=3 biological replicates. * p< 0.01, ** p< 0.001, *** p< 0.0001.

**Figure S4.**
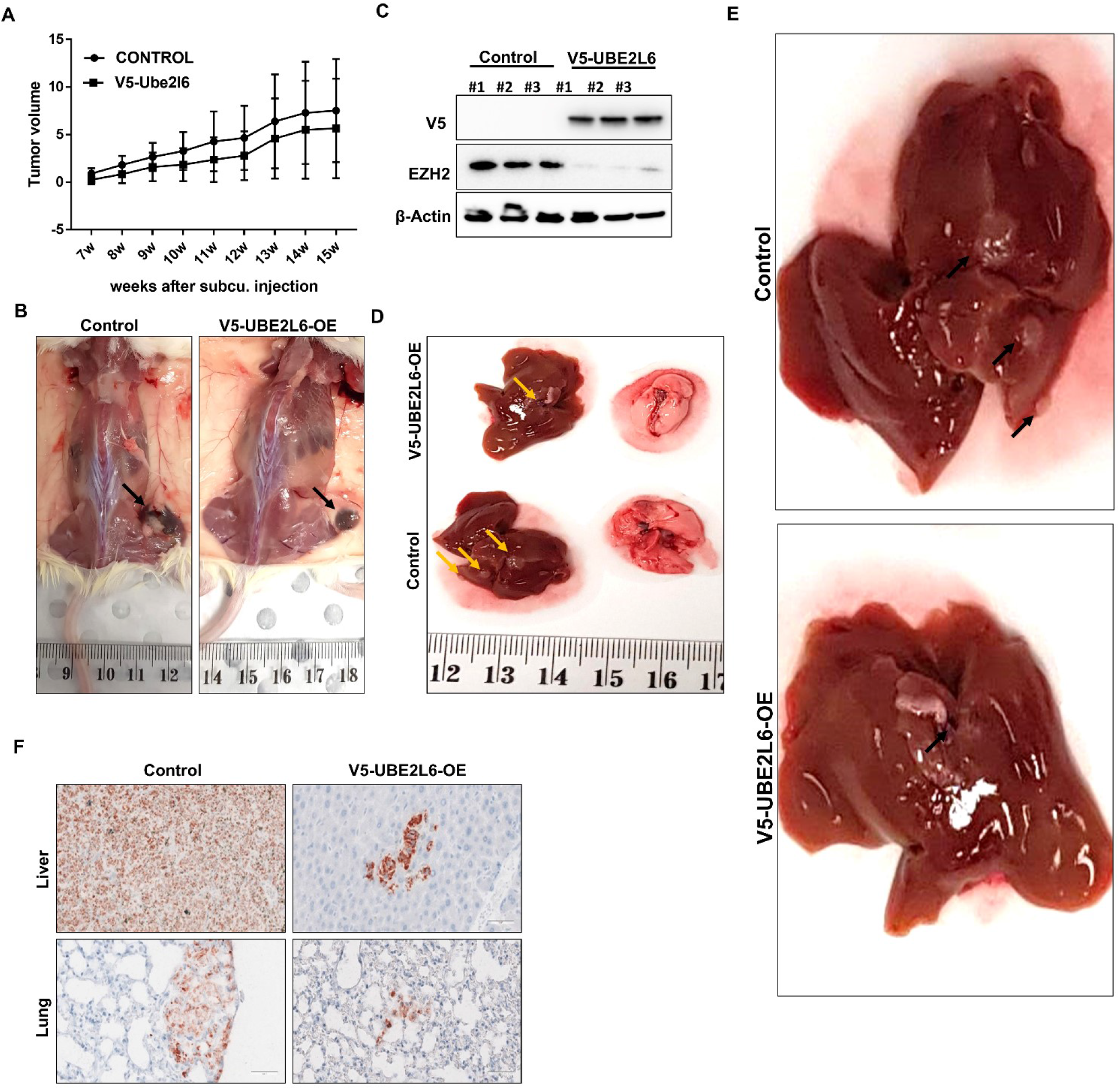
UBE2L6 overexpression decreases lung and liver metastasis rates *in vivo*. (A) Representative image of tumor bearing mice injected with LM-MEL28:B4:F3 harboring control or V5-UBE2l6-WT vector stably. (B) Western blot of three representative tumors with anti-V5 and anti-EZH2 antibody and (C) representative image of liver and lung from mice injected with B4:F3 harboring control or V5-UBE2l6-WT vector. (D) Higher magnification images of liver shown in (C). (E) Representative human mitochondria immunohistochemistry images (red) of lung and liver sections from mice that harboured a xenografted tumour that expressed either a control plasmid or V5-UBE2L6-WT.

**Figure S5.**
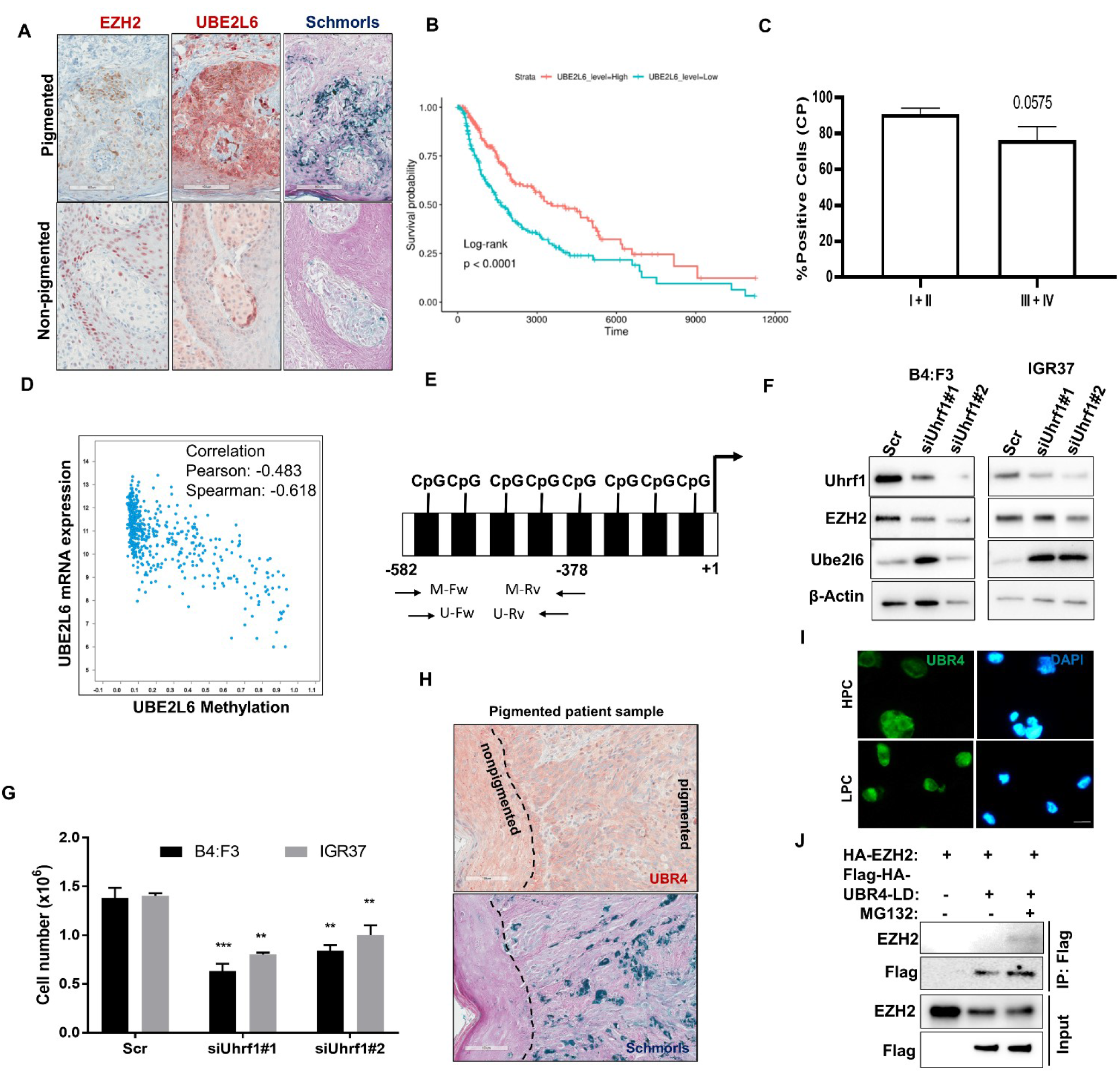
UHRF1 knockdown upregulates UBE2L6 and in turn downregulates EZH2 in melanoma cells. (A) Immunohistochemical staining of EZH2 and UBE2L6 in representative Schmorls stained pigmented and nonpigmented human melanoma specimens. Scale bar, 50 μm. (B) Kaplan-Meier survival of TCGA cutaneous melanoma patients (n = 427 patients), stratified by Ube2l6 mRNA levels. (C) Melanoma tumor staging in our human melanoma patient cohort stratified by cytoplasmic UBE2L6 protein levels. Protein score = the percentage of immune-positive cells. (D) Correlation between UBE2L6 mRNA and methylation levels in TCGA cutaneous melanoma samples. (E) Schematic representation of UBE2L6 promoter at CpG sites spanning from −582 to −378 relative to transcriptional start site (TSS). Forward primer (Fw) and reverse primer (Rv) used in methylation (m) and unmethylation (um) analysis are depicted. (F) UHRF1, EZH2 and UBE2L6 protein levels were determined with western blot and (G) viability with Trypan blue cell counting in B4:F3 and IGR37 cells transfected with scramble or two siUhrf1 oligos. (H) Immunohistochemical staining of UBR4 in representative Schmorls stained pigmented human melanoma specimens. Scale bar, 100 μm (I) UBR4 IF staining of HPC and LPC FACS-sorted from B16-F10 cells. (J) HA-tagged EZH2 and Flag-HA-tagged UBR-LD were coexpressed into HEK293 cells maintained in the absence or presence of MG132. The interaction between EZH2 and UBR4-LD was determined by immunoprecipitation with anti-Flag antibody followed by western blotting with anti-EZH2 antibody. n=3 biological replicates. ** p< 0.001, *** p< 0.0001.

